# Periweaning diet-induced activation of an IFNγ-mediated regulatory circuit promotes the homeostasis of cytotoxic CD8^+^ T Cells

**DOI:** 10.64898/2026.03.29.714815

**Authors:** Doğuş Altunöz, Ramin Shakiba, Kaushikk Ravi Rengarajan, Hamsa Narasimhan, Nikos E. Papaioannou, Sadiq Nasrah, Jessica Vetters, Maria L. Richter, Maria Parra Reyes, Nadine Nuschele, Denise Messerer, Sabine Schwamberger, Andreas Goschin, Dimitrios Starfas, Melanie Schmid, Tobias Straub, Michele Proietti, Katrin Böttcher, Maria Colomé-Tatché, Dirk Haller, Jan P. Böttcher, Stephanie Ganal-Vonarburg, Sophie Janssens, Christian Schulz, Anne B. Krug, Barbara U. Schraml

## Abstract

Balancing pathogen defence with maintaining tolerance to environmental antigens, such as food or commensals, in neonates is essential for survival and the establishment of life-long immune homeostasis. Instructed by environmental signals type 1 conventional dendritic cells (cDC1) drive either T cell tolerance or immunity. Here, we uncover an interferon (IFN)-γ-driven regulatory circuit in early life that relays dietary cues to spleen cDC1. Loss-of-function demonstrates that IFNγ-mediated STAT1-signaling induces an immunogenic maturation program in spleen cDC1 that instructs cDC1 to expand effector memory CD8⁺ T cells. This program emerges during weaning, when IFNγ production from lymphocytes rises, it occurs in germ-free mice and remains responsive to dietary intervention in adult mice. During the transition from breastfeeding to solid food at weaning, this circuit relays dietary information to spleen cDC1 to shape the effector phenotype of food-antigen specific CD8^+^ T cells in a feed-forward manner, allowing cDC1 to recalibrate the T cell pool at the moment of nutritional independence.

## INTRODUCTION

Early postnatal life is a critical period for developmental immune programming during which long-lasting immune phenotype and disease susceptibility are established^1–3^. During this time specific regulatory circuits ensure tolerogenic immune responses against environmental antigens, such as food and commensal microbes, that can suppress inflammatory events even in later life^4–6^. Simultaneously, immune cells are programmed with effector functions that are beneficial for anti-pathogen defense but can be harmful if directed against environmental antigens^7,8^. Understanding these immunostimulatory circuits holds great promise for treating allergy, improving vaccine efficacy or boosting immunity in diseases characterized by immunosuppression, such as cancer^1^.

Conventional dendritic cells (cDCs) are potent orchestrators of T cell mediated immunity^9,10^. Depending on the context in which they encounter antigen, such as the presence or absence of infection, inflammation, or tissue damage, cDCs can drive immunogenic or tolerogenic T cell responses^9,10^. cDCs exist as distinct subtypes with specific functions in pathogen defense. Type 1 conventional dendritic cells (cDC1) are potent activators of CD4^+^ T helper (Th) 1 and cytotoxic T cell responses against intracellular pathogens^9–12^. cDC2 on the other hand preferentially promote Th2 and Th17 responses against extracellular pathogens, as well as T follicular helper cell differentiation^9,10,13–16^. In addition to these well-defined roles in anti-pathogen defense both cDC1 and cDC2 can adopt functional states that promote T cell tolerance or activation in response to environmental cues that are often tissue specific^6,17–20^.

One mechanism through which cDCs acquire functional states is a process known as maturation^9,21,22^. Upon recognition of pathogens via pattern recognition receptors (PRRs) both cDC types upregulate expression of antigen presentation machinery, costimulatory molecules and C-C chemokine receptor type 7 (CCR7) - a process called “immunogenic” maturation in reference to stimulation by pathogenic signals^9,21,22^. CCR7-expressing cDCs can migrate to T cell areas of secondary lymphoid organs where they subsequently initiate T cell responses^9,21,23^. The maturation into CCR7^+^ cDCs also takes place in steady-state in the absence of microbial signals and in germ-free conditions and is thus referred to as “homeostatic” maturation^21,24^. Homeostatic cDC maturation is thought to primarily promote T cell tolerance^21^. In cDC1, for instance, the homeostatic maturation into CCR7-expressing cells can be induced by uptake of apoptotic cells and subsequent LXRβ-dependent cholesterol efflux^25^. Loss of LXRβ signaling in cDC1 causes increased activation of B and T cells in steady state adult mice^25^. In the thymus of adult mice, thymic epithelial cells produce type I and III IFNs (IFN-I and IFN-III) that promote the maturation of cDC1 into CCR7^+^ cells^26^. Whether this IFN-induced maturation involves the recognition of apoptotic cells is unclear but it occurs independent of microbial signals and is required for the thymic selection of regulatory T cells^26^. Thus, in homeostatic conditions cDCs integrate tissue-derived signals, including apoptotic cell death and cytokines, to promote T cell tolerance.

Early life is a period of substantial immune challenge during which the immune system must continuously interpret and respond to a rapidly changing environment. The most profound changes to the external environment are dietary changes during weaning and the initial encounter with commensals and pathogenic microbes^27,28^. Although early-life cDCs were once considered functionally immature to accommodate these challenges, accumulating evidence suggests they are instead finely tuned to their developmental context allowing them to establish and maintain homeostasis^3,6,29–33^. Apoptotic cell-driven cDC1 maturation is thought to suppress T cell responses to dying self during lung remodeling in neonatal mice^31^. In the pre-weaning period compared to adulthood, cDC2 exhibit a heightened ability to promote peripheral Treg differentiation in murine skin, lung, and spleen^6,29,30^. In neonatal skin cDC2 induce commensal-specific Tregs during a defined developmental window that prevent inflammation in later life^5,6^. The signals that drive the tolerogenic potential of cDC2 in neonatal skin and spleen are undefined, but in neonatal lung commensal encounter triggers a tolerogenic program in cDC2^29^. On the contrary, early microbial colonization enables spleen cDC1 to elicit Th1 responses to immunization with Ovalbumin (OVA) and protective CD8⁺ T cell responses to *Listeria monocytogenes* suggesting the induction of immunogenic programs by commensal encounter^32,33^. Similarly, in adult mice, commensal microbes drive IFN-I production from plasmacytoid DCs (pDCs) that transcriptionally poises splenic cDCs with inflammatory capacity^34^. Of note, transcriptional analyses show higher IFN-induced genes in spleen cDC2 of adult compared to young mice but lack of commensals does not cause cDC2 from adult mice to acquire a phenotype resembling cDC2 from young mice^30^. Thus, the functions of cDCs appear to be shaped by commensal microbiota in a tissue and age-specific manner.

Underscoring the influence of tissue-context on early-life cDC functions, neonatal cDC2 exhibit a Th2 bias in the lung, but not spleen, that is driven by type 2 cytokines during lung alveolarization^30,35^. In neonatal spleen both cDC1 and cDC2 are functionally competent to induce inflammatory T cell responses^30,32,36^, while in murine Peyer’s patches cDCs acquire immunocompetency only after weaning^37^. cDC immunocompetency at this site is reached independent of the microbiota and follows the differentiation of follicle-associated epithelium (FAE) M cells^37^. Thus, cDCs do not reach immunocompetency in all tissues simultaneously, suggesting tight regulation of cDC function across developmental age that is coordinated by tissue specific signals and external factors, such as commensals.

Here, we set out to define regulatory circuits that determine cDC1 function in spleen during early life and weaning. We find that the dietary switch at weaning triggers IFNγ production from lymphocytes independent of the microbiota. IFNγ pushes cDC1 in the spleen towards an immunogenic cell state that amplifies effector memory CD8^+^ T cells. This functional state of cDC1 arises in a STAT1-dependent manner, is characterised by CXCL9 expression and distinct from CCR7^+^ homeostatically matured cDC1. We find dietary cues are relayed to spleen cDC1 in a feed forward manner to shape the effector phenotype of food-antigen specific CD8^+^ T cells at distal sites. cDC1 maturation remained responsive to dietary intervention in adult mice, highlighting the therapeutic potential of dietary intervention to modulate dendritic cell function beyond the weaning period.

## RESULTS

### cDC1 from spleens of young and adult mice exhibit functional and transcriptional differences

We first confirmed that cDC1 from spleens of young and adult mice show distinct responsiveness to stimulation with pathogen-associated molecular patterns (PAMPs)^38,39^. We sort-purified XCR1^+^ cDC1 from spleens of 2-week-old and adult mice and measured cytokine production after stimulation with different PRR agonists (Fig. 1a-d, Supp. Fig. 1a). cDC1 from adult mice stimulated with the Toll-like-receptor (TLR) 2 and Dectin-1 ligand Zymosan, showed higher TNF production than those from 2-week-old mice (Fig. 1a). Similarly, depleted Zymosan, an exclusive Dectin-1 ligand, induced higher TNF, IL-23 and IL-1β production in cDC1 from adult compared to 2-week-old mice (Fig. 1b). In contrast, cDC1 from 2-week-old mice showed higher production of IL-10 and IL-12p70, the active form of IL-12, in response to CpG-B stimulation (Fig. 1c). Similarly, poly I:C stimulation induced stronger production of IL-12p40 and IL-6 in cDC1 from 2-week-old compared to adult mice (Fig. 1d). TNF production in response to CpG-B and poly I:C was similar across age (Fig. 1c, d). Thus, cDC1 from neonates and adult mice show qualitative differences in cytokine production after PRR stimulation as expected^39^.

**Figure 1:**
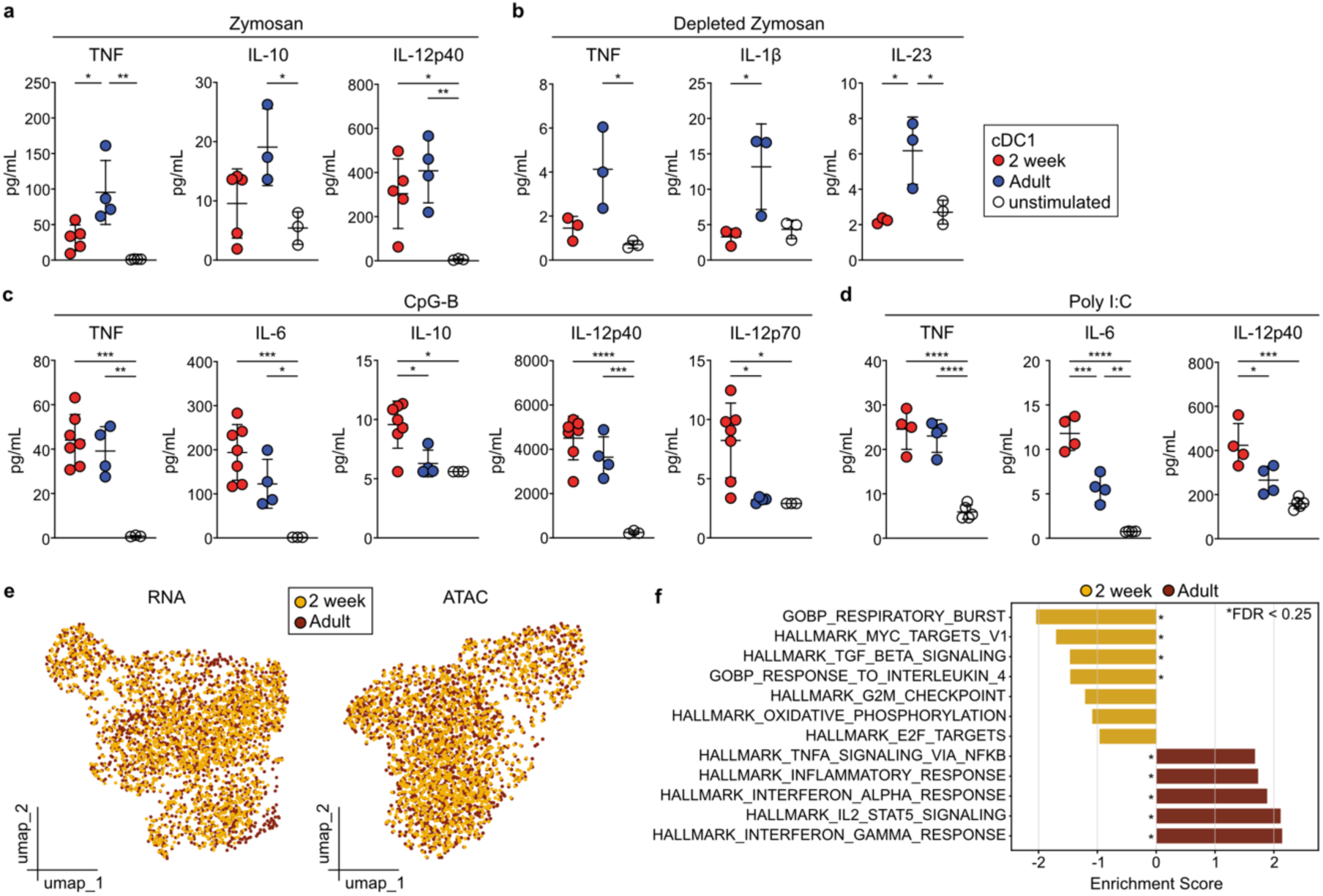
cDC1 from spleen of young and adult mice exhibit functional and transcriptional differences. **a-d,** Splenic cDC1 from 2-week-old or adult mice were left untreated or stimulated with Zymosan **(a)**, Depleted Zymosan **(b)**, CpG-B **(c)**, or Poly I:C **(d)**. After 20 hours cytokine concentrations in supernatant were quantified. Data points from unstimulated samples contain cDC1 from 2-week-old and adult mice. Each dot represents one biological replicate from two independent experiments (**a, c**). Data in b, d are representative of two independent experiments, horizontal bars represent mean, error bars represent SD. **e-f,** Cells annotated as cDC1 from 2-week-old and adult mice were isolated from a published dataset^40^. **e,** RNA-based and ATAC-based UMAP of splenic cDC1 with cells from 2-week-old and adult mice colored by age. **f**, GSEApy was run with default parameters and enrichment scores of some gene sets differentially regulated between cDC1 from 2-week-old and adult mice are shown. Statistical analysis was performed using one-way ANOVA with Tukey’s multiple comparisons test **(a-d).** Only statistically significant comparisons are indicated. *p < 0.05, **p < 0.01 ***p< 0.001 ****p < 0.0001.

Analysis of a single cell RNA and ATAC sequencing dataset of splenic cDCs from 2-week-old and adult mice^40^ revealed that the transcriptional identity of cDC1 was similar across age in uniform manifold projection (UMAP) space (Fig. 1e). Expression of *Tlr2, Tlr3* and *Tlr9* transcripts was low but similar across age (Supp. Fig 1b). Gene set enrichment analysis (GSEA), however, identified several pathways in XCR1^+^ cDC1 differentially regulated by age (Fig. 1f). cDC1 from adult mice were enriched for genes downstream of IFN-I and IFNγ (Fig. 1f), which is in line with our prior observations that spleen cDC2 from adult versus 2-week-old mice show enrichment of IFN-stimulated genes^30,40^.

### Spleen cDC1 transcriptionally change during weaning

Because commensal-induced IFNs transcriptionally shape spleen cDCs^34,41^ and weaning - the transition from breast milk to chow - correlates with a strong increase in microbial diversity^4^, we hypothesized that weaning might be a critical period in the functional regulation of splenic cDC1. To address this hypothesis, we performed single cell RNA-sequencing (scRNA-seq) of CD11c^+^MHCII^+^ splenocytes from mice aged 1, 3, 4 and 6 weeks (Fig. 2a, Supp. Fig. 2a). All mice were separated from their mothers at exactly three weeks of age and time points were chosen to cover the perinatal period around weaning until shortly before the onset of sexual maturity. After quality control we retained gene expression profiles from 7,689 cells with similar contribution from each time point (week 1: 2,131 cells, week 3: 2,050 cells, week 4: 1,752 cells, week 6: 1,756 cells). Unsupervised graph-based clustering of cells from all time points resulted in 15 clusters that segregated into two main metaclusters: cDC1 (clusters 0, 1, 4, 11, 12)^42^ and a cluster containing cDC2/DC3 (clusters 2, 5, 6, 7, 9)^42–44^, transitional DCs (tDCs; cluster 3)^45^ and CCR7^+^ migratory cDC (cluster 8)^46^ (Fig. 2b, Supp. Fig. 2b-f, Supp. Table 1). Cluster 14 could be identified as RORγt^+^ DCs^30,40^ (Supp. Fig. 2d, f), whereas cluster 13 was likely a contamination with *Gypa* expressing erythrocytes^47^ (Supp. Fig. 2d). Although we regressed cell cycle genes as cDCs proliferate in the developing spleen before weaning^30^, we found that clusters 10, 11 and 12 mainly consisted of proliferating cells (Supp. Fig. 2b, c). Homeostatically matured *Ccr7*^+^ cDC1 downregulate XCR1^24,25,48^ and in line with this we observed low expression of *Xcr1* in *Ccr7*^+^ cDCs (Supp. Fig. 2d). Of note, all cDC clusters contained cells from each time point, demonstrating that cDCs preserve their overall identity independent of developmental age (Supp. Fig. 2g).

**Figure 2:**
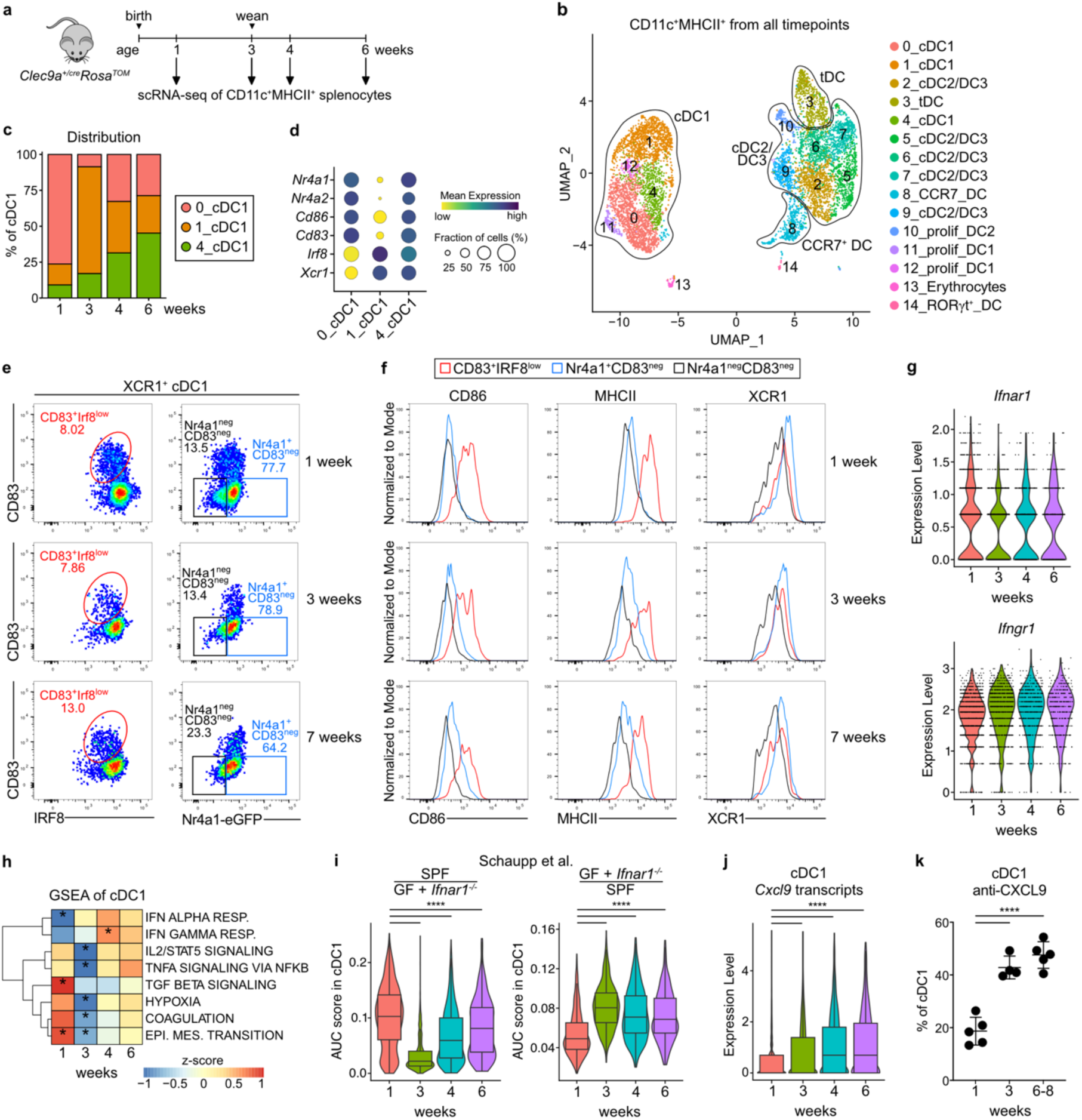
Spleen cDC1 transcriptionally change around weaning. **a,** Experimental set up. CD11c^+^MHCII^+^ cells were sorted from spleens of mice at the indicated ages and equal numbers from each timepoint were used as input for scRNA-seq. **b,** UMAP display of 7689 CD11c^+^MHCII^+^ cells grouped from all ages and annotated by cell type. **c,** Relative frequency of cDC1 clusters 0, 1 and 4 at the different time points. **d,** Bubble plot displaying expression of indicated genes across non-proliferating cDC1 clusters 0, 1 and 4. **e,** Flow cytometry of XCR1^+^ cDC1 from splenocytes from *Nr4a1^eGFP^* mice of indicated ages (for gating strategy see Supp. Fig. 2h). XCR1^+^ cDC1 were divided into CD83^+^IRF8^low^ (red), GFP^+^CD83^neg^ (blue) and GFP^neg^CD83^neg^ (black) populations. **f,** Histograms showing expression of indicated surface markers in CD83^+^IRF8^low^ (red), GFP^+^CD83^neg^ (blue) and GFP^neg^CD83^neg^ (black) cDC1. **g-k,** cDC1 clusters 0, 1 and 4 were grouped for analyses. **g,** Expression levels of *Ifnar1* and *Ifngr1* in cDC1 across age. **h,** Gene set enrichment analysis of cDC1 across age. Only gene sets with p < 0.01 for at least one time point are shown. **i,** AUC scores for genes described to be regulated by IFNAR in cDCs^34^. Boxes represent interquartile range, horizontal bars represent median, whiskers represent minimum and maximum. **j,** *Cxcl9* expression in cDC1 clusters (0, 1, 4) across age. Boxes represent interquartile range, horizontal bars represent median, whiskers represent minimum and maximum **(i,j)**. **k,** CXCL9 expression in XCR1^+^ cDC1 was quantified by flow cytometry in spleens from mice at the indicated ages. Each dot represents one biological replicate from two independent experiments. Statistical analysis was performed using multiple t-tests corrected for multiple comparisons using the Holm-Šídák method (**h-j**) or one-way ANOVA with Tukey’s multiple comparisons (**k**), *p < 0.05, **p < 0.01, ***p < 0.001, ****p < 0.0001.

Since we had shown that XCR1^+^ cDC1s exhibit age-dependent differences in cytokine production after PRR simulation (Fig. 1) we focused on *Xcr1*^+^ cDC1 clusters 0, 1, and 4 for further analyses. We noticed that the abundance of clusters 0, 1 and 4 within cDC1 changed with age and stabilized by 4 weeks of age (Fig. 2c). Cluster 0 showed lower expression of *Irf8* and *Xcr1* compared to clusters 1 and 4, while cluster 1 expressed lowest levels of *Nr4a1, Nr4a2, Cd86* and *Cd83* (Fig. 2d, Supp. Table 2), suggesting the clusters could be distinct maturation states. Indeed, flow cytometry of XCR1^+^ cDC1 from spleens of *Nr4a1^eGFP^* reporter mice confirmed that XCR1^+^ cDC1 could be divided into CD83^+^GFP^+^, CD83^neg^GFP^+^ and CD83^neg^GFP^neg^ cells (Fig. 2e, Supp. Fig. 2h). CD83^neg^GFP^neg^ cells showed lowest levels of CD86 and MHCII and most closely resembled cluster 1, while CD83^neg^GFP^+^ cells expressed intermediate levels of CD86 and MHCII and most closely resembled cluster 4 (Fig. 2d-f). The CD83^+^XCR1^+^ population most closely resembled cluster 0 with relatively low levels of IRF8 and highest levels of CD86 and MHCII at all ages (Fig. 2d-f). Thus, cDC1 exist as distinct maturation states, with CD83^+^ cells representing a more mature state of cDC1 than CD83^neg^ cells.

To gain insights into specific pathways that could regulate cDC1 maturation around weaning, we performed GSEA (Fig. 2h). We noticed that *Xcr1*^+^ cDC1 from 1-week-old mice showed lower expression of genes downstream of IFNα and IFNγ compared to *Xcr1*^+^ cDC1 from all other time points, although expression of *Ifnar1* and *Ifngr1* was similar across age (Fig. 2g, h). Accordingly, we found lower expression of genes induced by microbiota in an interferon-alpha/beta receptor (IFNAR) dependent manner^34^ and conversely higher expression of genes suppressed by IFNAR signaling^34^ in cDC1 from one compared to 3-week-old mice (Fig. 2i). Additionally, we found that the expression of the IFN-stimulated gene *Cxcl9*^49,50^ increased in cDC1 between one and 3 weeks of age, which was confirmed by protein staining (Fig. 2j, k). Together, these results identify IFN signaling as a potential driver of age-dependent functional programs in splenic cDC1 around weaning.

### IFNγ production from spleen lymphocytes rises with age

IFN-I, IFN-II and IFN-III stimulate the expression of similar genes. To start to address which type of IFN could be imprinting cDC1 with age, we profiled the expression of IFN-I (IFNα, IFNβ), IFN-II (IFNγ) and IFN-III (IFNλ2/3) in the spleen using quantitative real time PCR (qRT-PCR; Fig. 3a). We observed that the expression of *Ifna4* and *Ifnl2/3* was constant in spleens of mice across age (1, 2, 3, 4 and 10 weeks; Fig. 3a). In contrast, expression of *Ifng* increased steadily from one to 4 weeks of age and somewhat declined thereafter (Fig. 3a).

**Figure 3:**
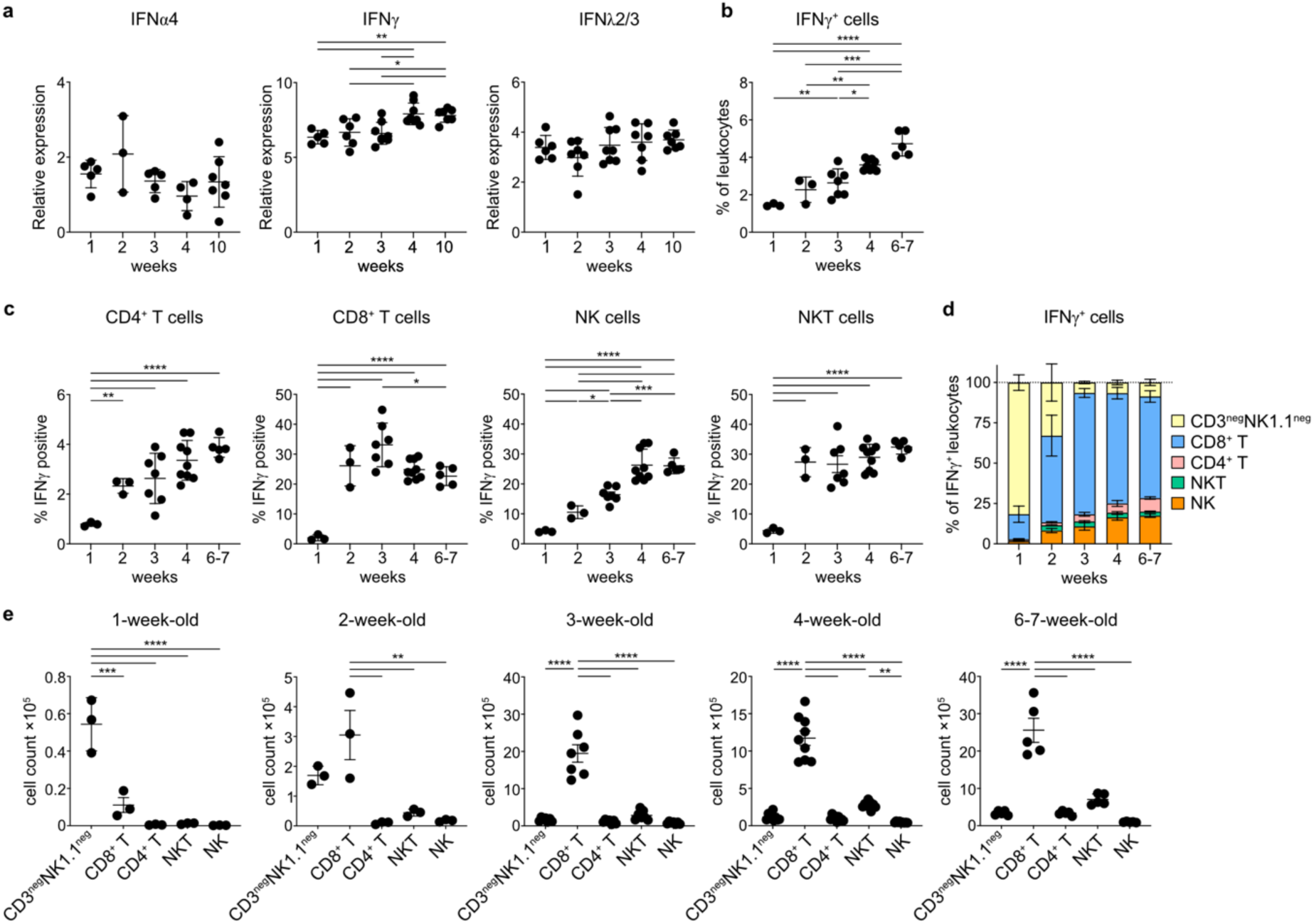
IFNγ production from spleen lymphocytes rises with age. **a,** Quantification of *Ifna4*, *Ifng* and *Ifnl2/3* expression in total spleen tissue by quantitative real time PCR (qRT-PCR). Expression relative to *Actb* is shown. **b-e,** Splenocytes from mice at the indicated ages were cultured for 5 hours with phorbol 12-myristate 13-acetate (PMA) and ionomycin. Brefeldin A was added for the last 3 hours. Cells were then stained for surface markers and IFNγ and analyzed by flow cytometry. **b,** Frequency of IFNγ-producing leukocytes across age. **c,** Frequency of IFNγ-producing CD4^+^, CD8^+^, NKT and NK cells at the indicated ages. **d,** Contribution of different lymphocyte subsets to total IFNγ^+^ cells. **e,** Cell numbers of IFNγ-producing leukocytes across age. Each dot represents one biological replicate pooled from at least 2 independent experiments, horizontal bars represent mean, error bars represent SD. Statistical analysis was performed using one-way ANOVA with Tukey’s multiple comparisons test. Only statistically significant comparisons are indicated, *p < 0.05, **p < 0.01 ***p< 0.001 ****p < 0.0001.

Accordingly, we observed that the frequency of IFNγ producing lymphocytes in spleen increased steadily from one week to 7 weeks of age (Fig. 3b, Supp. Fig. 3a). IFNγ production increased with age in NK and NKT cells, as well as in CD4^+^ T cells (Fig. 3c) but spiked at week three in CD8^+^ T cells (Fig. 3c). Notably, CD8^+^ T cells constituted the biggest fraction of IFNγ-producing cells at all time points except one week of age, when most IFNγ was produced by CD3^neg^NK1.1^neg^ cells (Fig. 3d, e).

### IFNγ mediates STAT1-dependent CXCL9 production in cDC1 independent of the microbiota

Signaling downstream of all types of IFN is mediated by the transcription factor signal transducer and activator of transcription 1 (STAT1)^51,52^. We therefore crossed *Stat1^flox^* mice to *Itgax^cre^* and *Clec9a^cre^* mice^53,54^ to achieve deletion of IFN-responsiveness specifically in cDCs. At 3 weeks of age, when we had observed the strongest IFN-induced gene expression (Fig. 2h), we profiled cDC1 from both *Itgax^cre^Stat1^fl/fl^*and *Clec9a^+/cre^Stat1^fl/fl^* mice for CXCL9 production. CXCL9 was reduced in both the frequency and the amount of CXCL9 produced per cell in cDC1 from *Itgax^cre^Stat1^fl/fl^* and *Clec9a^+/cre^Stat1^fl/fl^* compared to control mice, however, CXCL9 production was not completely abolished (Fig. 4a, b, Supp. Fig 4a,b). Of note, STAT1-deficient cDC1 also showed reduced production of IL-12p40 (Fig. 4a, b, Supp. Fig 4a,b), a marker of homeostatic cDC maturation^55^. Interestingly, we observed no differences in CXCL9 or IL-12p40 production from cDC1 from 3-week-old mice with constitutive deletion of *Ifnar1* (Supp. Fig. 4c) or with deletion of *Ifnar1* specifically in cDCs (*Clec9a^+/cre^Ifnar1^fl/fl^* mice, Fig. 4c). In contrast, cDC1 from 3-week-old mice lacking IFNγ receptor (*Ifngr1^-/-^*) showed reduced CXCL9 production compared to age-matched *Ifngr1^+/-^* controls, whereas IL-12p40 production was unaltered between genotypes (Fig. 4d). Of note, among cDC subsets, cDC1 were the dominant producers of CXCL9, whereas production from cDC2 or pDCs was negligible (Supp. Fig. 4d). Thus, CXCL9 production in cDC1 from weanling mice is regulated by IFNγ in a STAT1-dependent manner.

**Figure 4:**
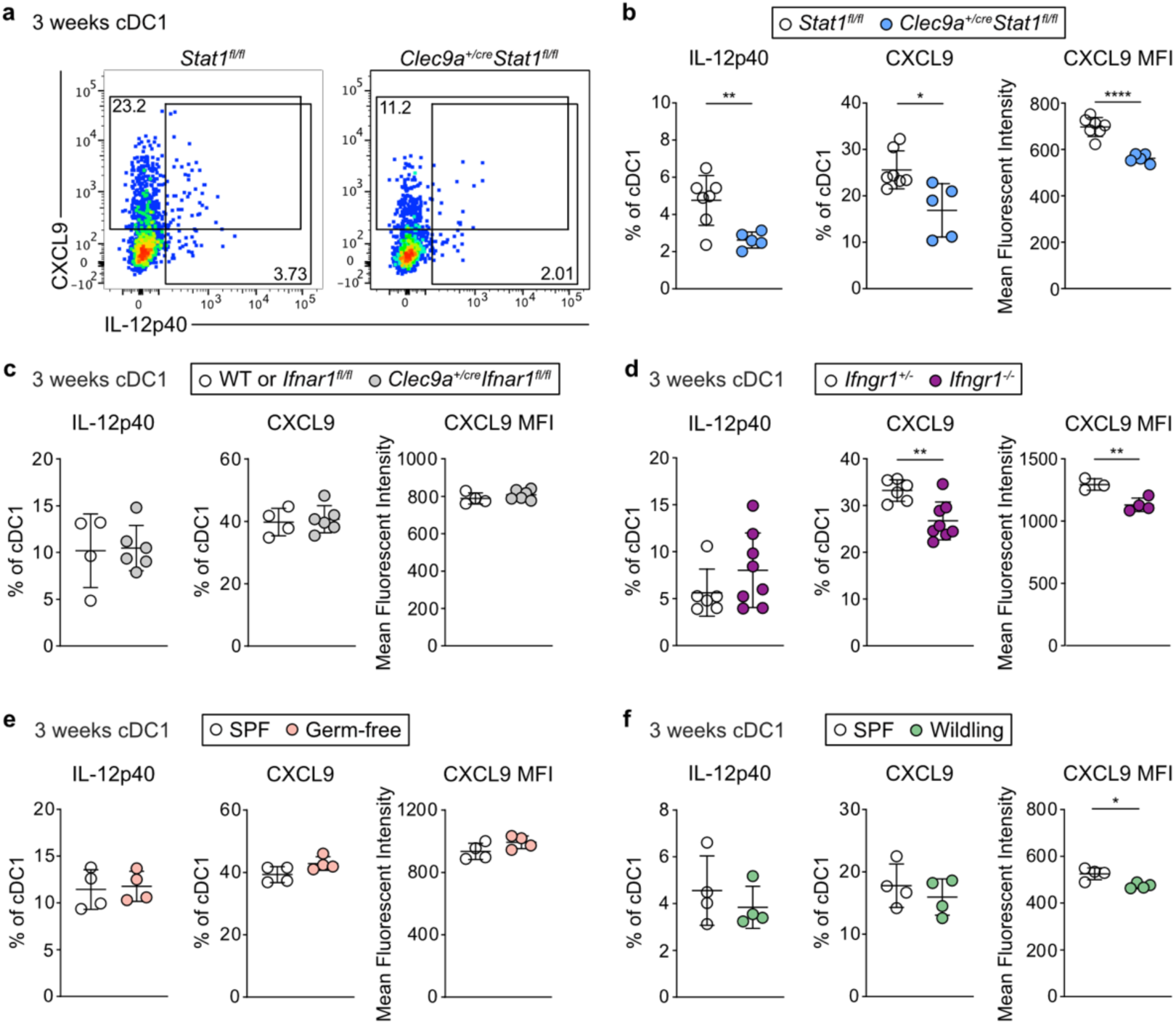
IFNγ mediates STAT1-dependent CXCL9 production in cDC1 independent of the microbiota. **a-f** Splenocytes from 3-week-old mice of the indicated genotypes were cultured for 4 hours with Brefeldin A and Monensin. XCR1^+^ cDC1 were then analyzed for cytokine production by flow cytometry. **a,** CXCL9 and IL-12p40 staining profiles and **b** quantification in cDC1 from *Clec9a^+/cre^Stat1^fl/fl^* and *Stat1^fl/fl^* littermate controls. **c,** Quantification of IL-12p40^+^ and CXCL9^+^ cDC1 from 3-week-old *Clec9a^+/cre^Ifnar1^fl/fl^* and *Ifnar1^fl/fl^* littermates or age-matched wild type controls. **d,** Quantification of IL-12p40^+^ and CXCL9^+^ cDC1 from 3-week-old and *Ifngr1^+/-^*or *Ifngr1^-/-^* mice. **e-f,** cDC1 from 3-week-old GF and age-matched SPF mice **(e)** or 3-week-old wildling and age-matched SPF mice **(f)** were analyzed as above and IL-12p40^+^ and CXCL9^+^ cDC1 were quantified. Each dot represents one biological replicate pooled from two independent experiments (**a-d**). Horizontal bars represent mean, error bars represent SD. Statistical analysis was performed using two-tailed Welch’s *t*-test, *p < 0.05, **p < 0.01 ***p< 0.001 ****p < 0.0001.

Weaning demarcates the change from breastfeeding to grain-based chow diet in mice, which correlates with a strong increase in commensal microbial diversity^4^. Since commensal microbiota have been linked to IFN-I and IFNγ-mediated signaling pathways in spleen immune cells^34,41^, we assessed CXCL9 production from cDC1 in germ-free (GF) and wildling mice at weaning. GF mice are raised sterile and are not colonized by commensals^56^, while “wildling” mice harbor a highly diverse microbiome resembling that of mice in their natural habitat^57^. Notably, we observed no differences in CXCL9 or IL-12p40 production between cDC1 from age-matched mice housed under specific pathogen free (SPF) and GF conditions (Fig. 4e) or between wildling and SPF mice (Fig. 4f).

### STAT1-signaling in cDC1 affects the homeostasis of CD8^+^ T cells in murine spleen

The above data show that IFNγ induces CXCL9 production in a fraction of cDC1 independent of microbial stimuli, indicating that this constitutes a homeostatic maturation process. Since homeostatic cDC maturation promotes T cell tolerance^21,24–26^, we profiled splenic T cells in *Itgax^cre^Stat1^fl/fl^* and *Clec9a^+/cre^Stat1^fl/fl^* mice at 3-weeks-old. CD4^+^ and CD8^+^ T cells were distinguished into cells with a naïve (CD44^neg^CD62L^+^), central memory (T_CM_; CD44^+^CD62L^+^) and effector memory (T_EM_; CD44^+^CD62L^neg^) phenotype (Fig. 5a,b, Supp. Fig. 5a-c). Additionally, we profiled Foxp3^+^ regulatory T cells. We found no consistent differences between genotypes in CD4^+^ T_EM_ or T_CM_ cells, as well as FOXP3^+^ Tregs (Supp. Fig. 5a-f). In contrast, within CD8^+^ T cells we observed a specific reduction of T_EM_ phenotype cells in *Itgax^cre^Stat1^fl/fl^*and *Clec9a^+/cre^Stat1^fl/fl^* mice compared to age-matched controls (Fig. 5a-b, Supp. Fig. 5g-h). CD8^+^ T cells with a T_CM_ phenotype were similar between genotypes (Fig. 5a-b, Supp. Fig. 5g-h).

**Figure 5:**
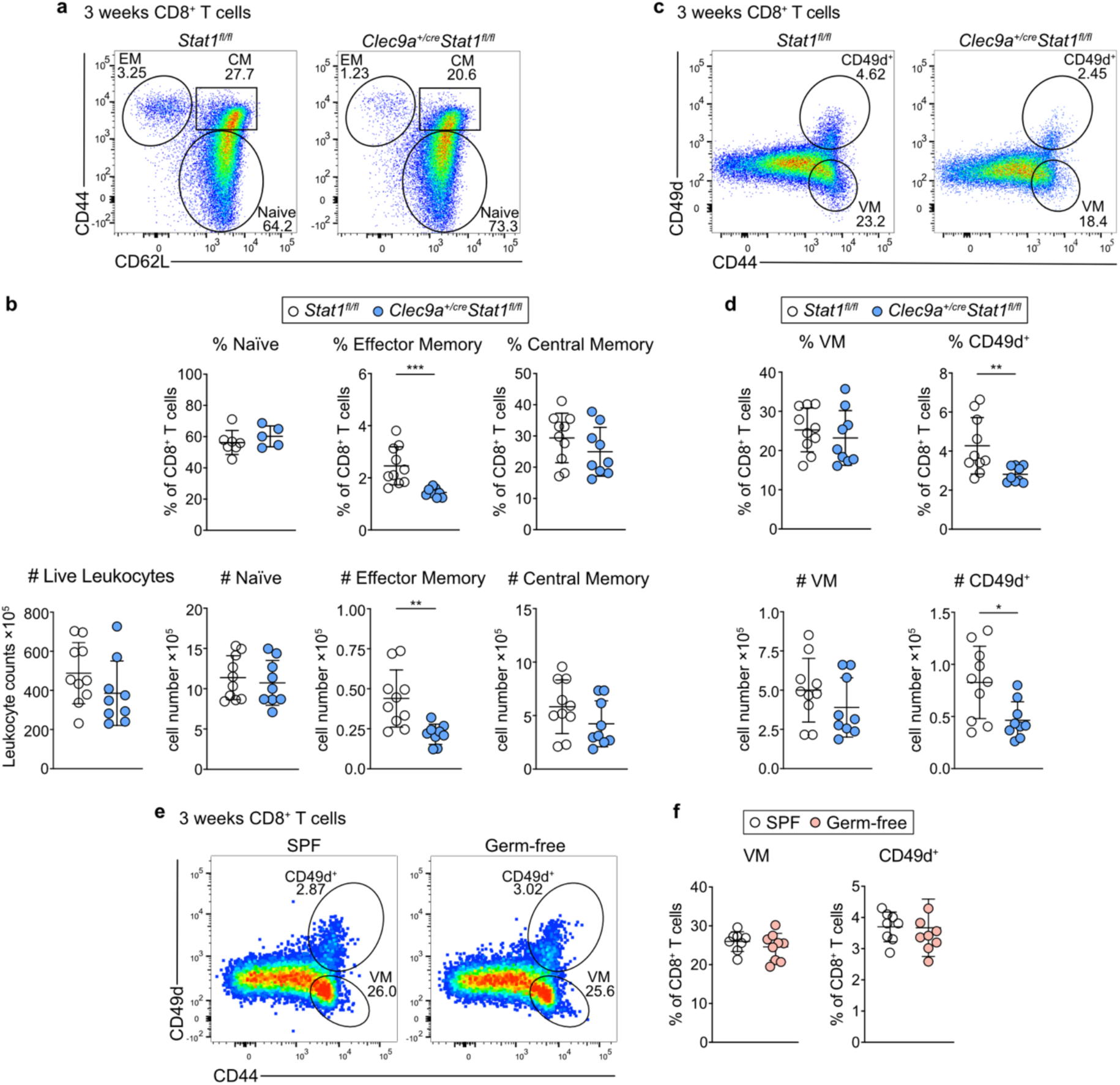
STAT1-signaling in cDC1 affects the homeostasis of CD8^+^ T cells in murine spleen. **a-b,** CD8^+^ T cells in the spleen were divided into effector memory (CD44^+^CD62L^neg^), central memory (CD44^+^CD62L^+^) and naïve (CD44^neg^CD62L^+^) populations. **a,** Representative gating and **b,** quantification of CD8^+^ T cell subsets in spleens of 3-week-old *Clec9a^+/cre^Stat1^fl/fl^* and *Stat1^fl/fl^* littermate controls. **c-d,** CD8^+^ T cells in the spleen were divided into CD49d^+^CD44^+^ antigen-experienced and CD49d^neg^CD44^+^ virtual memory (VM) populations. Gating (**c**) and quantification (**d**) of CD49d^+^ and VM populations in 3-week-old *Clec9a^+/cre^Stat1^fl/fl^* mice and *Stat1^fl/fl^* littermate controls. **e-f,** Representative gating (**e**) and quantification (**f**) of CD49d^+^ and VM populations in 3-week-old GF and age-matched SPF mice. Each dot represents one mouse pooled from at least two independent experiments, horizontal bars represent mean, error bars represent SD. Statistical analysis was performed using two-tailed Welch’s *t*-test, *p < 0.05, **p < 0.01, ***p < 0.001, ****p < 0.0001.

Although memory CD8^+^ T cells typically arise in an antigen-dependent manner in response to infection, memory-phenotype CD8^+^ T cells also exist in substantial numbers within hosts that have not been exposed to pathogens^58–60^. These memory-phenotype T cells entail so called ‘innate memory’ or ‘virtual memory’ T cells that are antigen-inexperienced and are propagated in the periphery by IL-15 and IL-4 produced by cDC1s^60–62^. They can be distinguished from antigen-experienced T cells by lack of integrin α4 (CD49d)^59,60,63,64^. How these antigen-experienced memory phenotype T cells arise in hosts that have not been exposed to pathogens and what antigens they recognize is unclear. We found a specific reduction of CD49d^+^CD44^+^CD8^+^ T cells in 3-week-old *Itgax^cre^Stat1^fl/fl^* and *Clec9a^+/cre^Stat1^fl/fl^* compared to control mice, whereas CD49d^neg^CD44^+^CD8^+^ virtual memory T cells were similar between genotypes (Fig. 5c, d, Supp. Fig. 5i). Of note, CD49d^+^ CD8^+^ T cells included cells with a T_EM_ and T_CM_ phenotype, however, only cells with a T_EM_ phenotype were reduced in *Clec9a^+/cre^Stat1^fl/fl^*compared to control mice (Supp. Fig. 5j, k). At 3 weeks of age these CD49d^+^CD44^+^ T_EM_ cells contained only few MR1-5-OP-RU tetramer positive circulating

Mucosal-Associated Invariant T (MAIT) cells, which also express CD8α, CD44 and CD49d and can be found in mouse spleen^65^ (Supp. Fig. 5l, m). Furthermore, the frequency of CD49d^+^CD44^+^CD8^+^ T_EM_ cells was similar between SPF and GF mice (Fig. 5e, f), further supporting that these cells are not MAIT cells, which are absent in GF mice^66^. Thus, antigen-experienced CD8^+^ T cells with T_EM_ phenotype that arise independent of microbial exposure are reduced in spleen during weaning in mice with cDC-intrinsic loss of *Stat1*.

### STAT1-signaling induces a specific immunogenic state of cDC1

To gain further mechanistic insights into the dysregulation of cDC1 and CD8^+^ T cell homeostasis in *Clec9a^+/cre^Stat1^fl/fl^* mice we performed scRNA-seq of CD90.2^+^ cells (including T cells, NKT cells and innate lymphocytes) and CD11c^+^MHCII^+^ cDCs from spleens of 3-week-old *Clec9a^+/cre^Stat1^fl/fl^*mice and *Stat1^fl/fl^* littermate controls (Supp. Fig. 6a). Unsupervised clustering of CD11c^+^MHCII^+^ cells from both genotypes resulted in 23 clusters, which were assigned as cDC1 (clusters 0, 3, 5, 8, 9, 11, 12, 20, 22), cDC2/DC3 (clusters 1, 2, 6, 7, 10, 14, 16, 17, 18, 19) and tDC (cluster 4) based on published gene signatures (Fig. 6a, Supp. Fig. 6b-d, Supp. Table 1). The resolution for unsupervised clustering was chosen to reveal *Sirpa-*expressing *Ccr7*^+^ cDC2 (cluster 13) and *Irf8*-expressing *Ccr7*^+^ cDC1^25^ (cluster 15, Supp. Fig. 6b, c). Again, we identified a small contaminating cluster of *Gypa*-expressing erythrocytes (cluster 21)^47^ which was excluded from further analyses (Supp. Fig. 6b).

**Figure 6:**
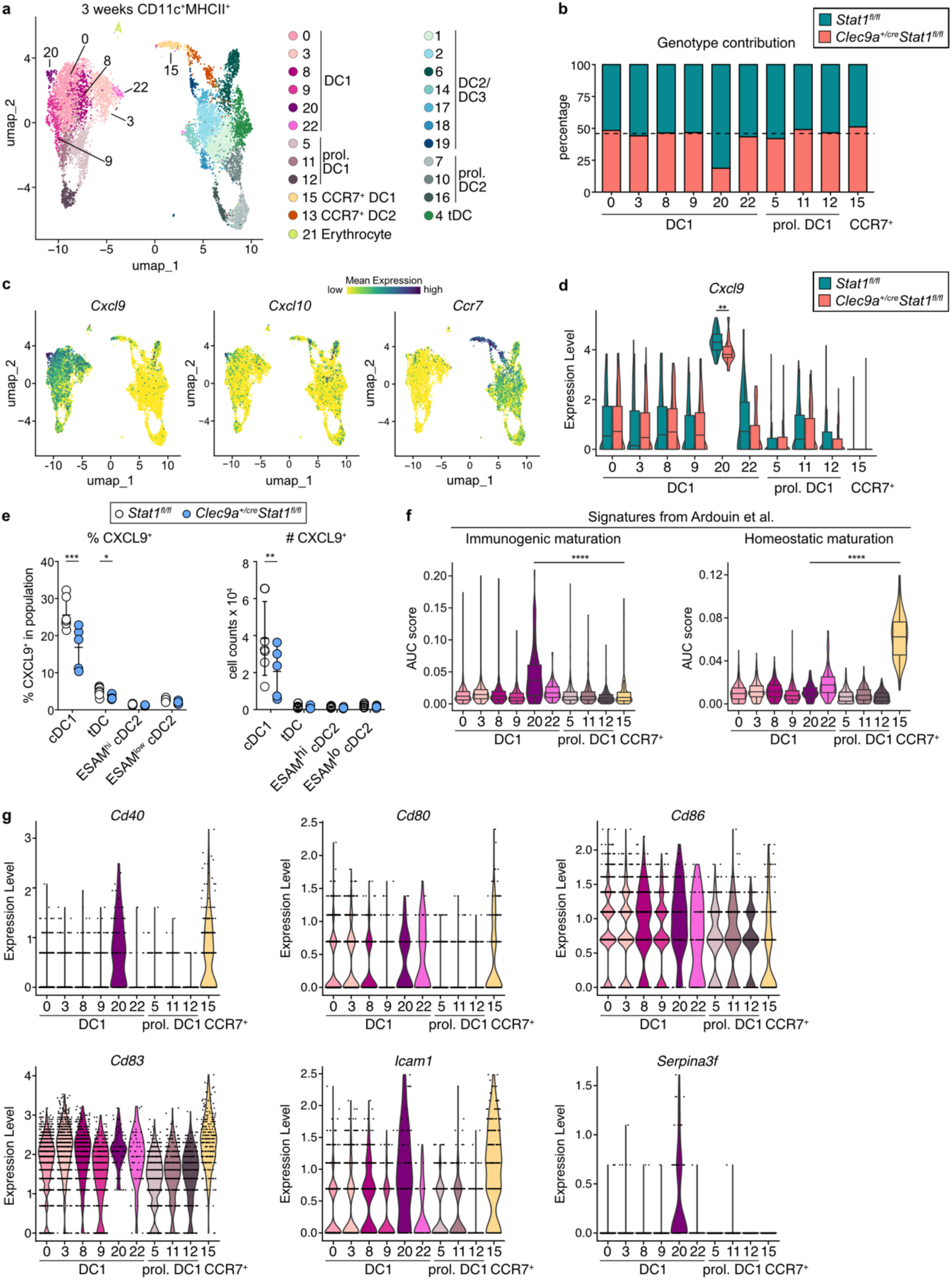
STAT1-signaling is required for an immunogenic state of cDC1. **a-d,** scRNA-seq of CD11c^+^MHCII^+^ cells from spleens of 3-week-old *Clec9a^+/cre^Stat1^fl/fl^* and *Stat1^fl/fl^*mice. **a,** UMAP display of 9906 CD11c^+^MHCII^+^ cells from both genotypes annotated by cell type (*Clec9a^+/cre^Stat1^fl/fl^*: 4935 cells, *Stat1^fl/f^*: 4971 cells). **b,** Contribution of cells from each genotype to individual cDC1 clusters. Dotted line is the contribution of cDC1 of *Clec9a^+/cre^Stat1^fl/fl^* genotype to all cDC1 clusters combined. **c,** Expression of *Cxcl9, Cxcl10,* and *Ccr7* projected on the UMAP display independent of genotype. **d,** Expression levels of *Cxcl9* in cDC1 clusters split by genotype. **e**, Splenocytes from 3-week-old mice were cultured for 4 hours with Brefeldin A and Monensin. cDC subsets were then analyzed for CXCL9 production by flow cytometry. Each dot represents one mouse pooled from at least two independent experiments, horizontal bars represent mean, error bars represent SD. **f,** AUC scores for immunogenic maturation or tolerogenic maturation signatures^24^ within cDC1 clusters. **g,** Expression of indicated genes in cDC1 clusters, irrespective of genotype. Box represents interquartile range, horizontal bar represents median, whiskers represent minimum and maximum enrichment scores (**f**), or gene expression level across cDC1 clusters (**d**, **g**). Statistical analysis was performed using unpaired t-test, **p < 0.01, ****p < 0.0001. For **f**, statistical testing was only performed between clusters 15 and 20.

Cells from both genotypes contributed equally to individual clusters assigned as cDC2/DC3 and tDC (Supp. Fig. 6e), while within cDC1, cluster 20 was dominated by cells from *Stat1^fl/fl^* controls (Fig. 6b). As expected, *Stat1* transcripts were reduced across cDC clusters from *Clec9a^+/cre^Stat1^fl/fl^*mice confirming efficient *Stat1* deletion (Supp. Fig. 6g). cDC1 cluster 20 expressed highest levels of *Cxcl9* and *Cxcl10* compared to other cDC1 clusters and lacked *Ccr7* (Fig. 6c, Supp. Table 3). Cluster 20 transcriptionally differed from the homeostatically matured *Ccr7*^+^ cDC1 cluster 15, which highly expressed *Il12b* and *Cd200* (Fig. 6c, d, Supp. Fig 6c, Supp. Table 3). In an intravenous labeling approach CCR7^+^ DCs poorly labelled (Supp. Fig. 6f), consistent with their localization in the white pulp^25^. In contrast, CXCL9^+^ DCs strongly labelled, indicating localization in blood exposed spleen regions, like red pulp or marginal zone (Supp. Fig. 6f). *Cxcl9* expression in cluster 20 was lower in cells from *Clec9a^+/cre^Stat1^fl/fl^* mice compared to control mice (Fig. 6d). Thus, not only fewer of the *Stat1*-deficient cDC1 acquire this transcriptional state but the ones that do express lower levels of *Cxcl9* (Fig. 6d). Of note, specifically cDC1, but not cDC2 or tDCs, showed reduced CXCL9 production in the absence of STAT1 (Fig. 6e).

cDC1 cluster 20 showed high *Stat1* expression and exhibited an enrichment of IFNα and IFNγ response genes (Supp. Fig. 6g, h), indicating it represents a distinct transcriptional state of cDC1 shaped by IFN signaling. We therefore reclustered the cDC1 and migratory DC clusters from the scRNA-seq dataset across age at higher resolution and also found a disctinct state marked by high levels of *Cxcl9* and *Cxcl10* (new cluster 9) that had previously been included in cluster 1 (Fig. 2c, Supp. Fig. 7a-d). This new cluster 9 increased drastically in frequency between one and three weeks of age (Supp. Fig. 7e, f).

PRR-induced immunogenic cDC1 maturation is distinguished from homeostatic cDC1 maturation by expression of specific genes^24^. Among cDC1 clusters, *Ccr7*^+^ cDC1 (cluster 15) from *Clec9a^+/cre^Stat1^fl/fl^* and *Stat1^fl/fl^* mice showed highest expression of genes associated with homeostatic cDC maturation^24^, whereas cDC1 cluster 20 showed highest expression of genes associated with immunogenic cDC maturation (Fig. 6f). Further, cluster 20 expressed a variety of co-stimulatory molecules indicative of mature antigen presenting cells, including *Cd80*, *Cd86*, *Cd83* and *Cd40*, as well as *Icam1*, which stabilizes the immunological synapse^67^ (Fig. 6g). Notably, Cluster 20 also expressed highest levels of the IFN-responsive cytosolic serpin *Serpina3f* (Fig. 6g) that is part of a family of proteins that protects from cell death, including cytotoxic T cell-induced killing^68,69^. Taken together these data show that STAT1-signaling in cDC1 controls the emergence of a unique transcriptional maturation state of cDC1.

### Loss of *Stat1* in cDCs impairs the communication between cDC1 and cytotoxic CD8^+^ T cells

We next performed unsupervised clustering of the CD90.2^+^ cells (Fig. 7a). This resulted in 20 clusters (Fig. 7a), two of which were excluded as monocyte contamination because they lacked *Cd3e* and *Thy1* and expressed typical monocyte markers (*Lyz2*, *Adgre4*) and genes associated with antigen presentation (Supp. Table 4). As expected, the remaining 18 clusters could be identified as *Klrb1c^+^* NK cells (clusters 5, 7, 16), *Klra1*^+^, *Klra7*^+^, *Cd3e*^+^ NKT cells (Clusters 4, 11), and *Cd3e* expressing CD4^+^ and CD8^+^ T cells (Fig. 7a, Supp. Fig. 8a). Of these, clusters 10, 13 and 12 contained mostly proliferating cells (Supp. Fig. 8b). Cells from both genotypes contributed equally to individual clusters except for CD8^+^ T cell clusters 14 and 17, which were predominated by cells from *Stat1^fl/fl^*control mice (Fig. 7b, Supp. Fig. 8c). Importantly, *Stat1* expression was similar between cells from *Clec9a^+/cre^Stat1^fl/fl^*and *Stat1^fl/fl^* control mice, confirming that *Clec9a^cre^* does not delete *Stat1* in CD90^+^ lymphocytes and any effects observed in T cells from *Clec9a^+/cre^Stat1^fl/fl^* mice are secondary to deletion of *Stat1* in cDCs (Supp. Fig. 8d). Clusters 0 and 6 most closely resembled naïve T cells (Supp. Fig. 8e). Cluster 15 showed highest expression of *Eomes* suggesting these cells constituted virtual memory T cells^60^ (Supp. Fig. 8e). Notably, cluster 17 expressed highest levels of *Stat1* and (Supp. Fig. 8d) and showed evidence of IFN-regulated genes (Supp. Table 4). Expression of *Sell* (encoding CD62L), *Cd44*, *Itga4* (encoding CD49d) and *S100a4*, identified cluster 14 as CD49d^+^ CD8^+^ T_EM_ cells, which, among CD8^+^ T cell clusters, showed highest expression of *Cxcr3*, the receptor for CXCL9, - 10, and – 11 (Fig. 7c, Supp. Fig. 8e). Cell-cell communication analysis between cDC and T cell clusters using *Community*^70^ revealed that the sender-receiver pairs 20 cDC1/14 CD8^+^ T (and reverse) as well as 20 cDC1/17 CD8^+^ T showed strong reduction in the number and weight of predicated interactions in *Clec9a^+/cre^Stat1^fl/fl^* mice compared to control mice (Fig. 7e). The expression of *Ccr7* on CD8^+^ T cell cluster 17, but not on cDC1 cluster 20, suggested that these cells reside in distinct regions of the spleen, leading us to focus on the interactions between cluster 20 cDC1 and cluster 14 CD8^+^ T cells. Community predicted CXCR3 and ICAM1-mediated interactions between these clusters (Fig. 7f). Cluster 14 further expressed high levels of *Gzmb*, *Gzmk*, *Ccl5* and *Cx3cr1*, markers for cytotoxic CD8^+^ T_EM_ cells^71,72^ (Figure 7d, Supp. Fig. 8e). Indeed, after stimulation sorted CD49d^+^CD44^+^ CD8^+^ T cells stained strongest for Granzyme B and the degranulation marker CD107a compared to virtual memory and naïve CD8^+^ T cells, confirming their effector phenotype (Fig. 7g, h). Of note, CD49d^+^CD44^+^ T_EM_ cells from *Clec9a^+/cre^Stat1^fl/fl^* mice showed comparable or slightly lower expression of Granzyme B comparted to T_EM_ cells from *Clec9a^+/+^Stat1^fl/fl^* control mice (Fig. 7i), indicating that the few CD49d^+^CD44^+^ T_EM_ cells that remain in *Clec9a^+/cre^Stat1^fl/fl^*mice possess effector function. Thus, loss of *Stat1* in cDCs specifically impairs the communication of a CXCL9-expressing state of cDC1 with CD8^+^ effector memory T cells.

**Figure 7:**
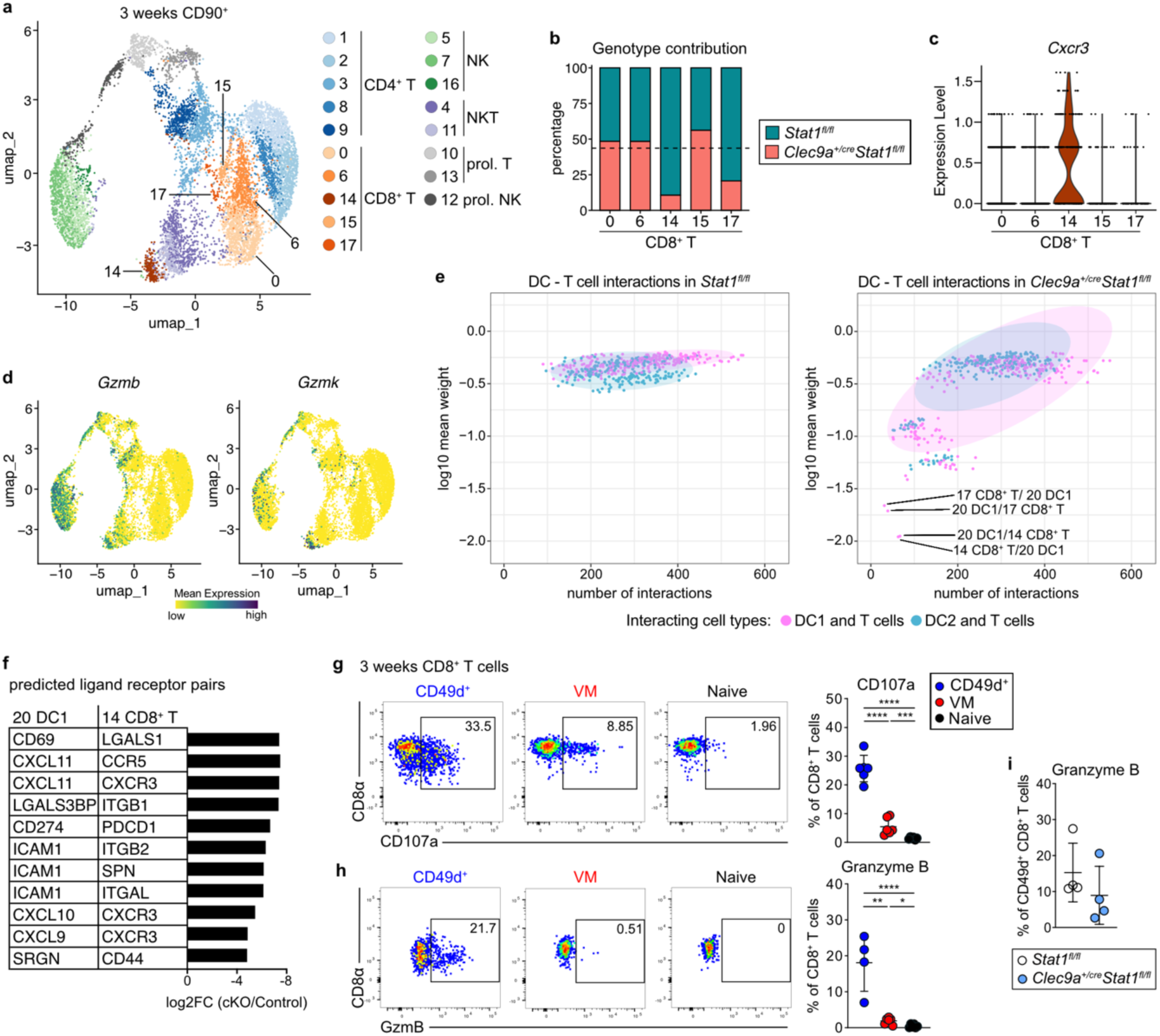
Loss of STAT1-signaling in cDCs impairs the communication between cDC1 and cytotoxic T cells. **a-d,** CD90^+^ cells were sort-purified from the spleens of 3-week-old *Clec9a^+/cre^Stat1^fl/fl^*and *Stat1^fl/fl^* mice and subjected to scRNA-seq. **a,** UMAP display of 8995 CD90^+^ cells from both genotypes annotated by cell type (*Clec9a^+/cre^Stat1^fl/fl^*: 4393, *Stat1^fl/f^*: 4602). **b,** Contribution of cells from each genotype to individual CD8^+^ T cell clusters. Dotted line represents the contribution of CD8^+^ T cells of *Clec9a^+/cre^Stat1^fl/fl^* genotype to all CD8^+^ T cell clusters combined. **c,** Expression levels of *Cxcr3* among CD8^+^ T cell clusters. **d,** Expression of *Gzmb, Gzmk* projected on the UMAP independent of genotype. **e-f,** Interactome analysis between cDC and T cell clusters was performed with *Community*. **e**, Number of interactions and average interaction intensity between pairs of DC – T cell clusters from *Stat1^fl/fl^* and *Clec9a^+/cre^Stat1^fl/fl^* mice is shown. **f**, Predicted ligand-receptor pairs between cluster 20 cDC1 and cluster 14 CD8^+^ T and respective log2FC of interaction intensity in *Clec9a^+/cre^Stat1^fl/fl^* over *Stat1^fl/fl^* control are shown. Pairs were filtered for interactions including ligands or receptors that showed differential gene expression in cluster 20 cDC1 vs. other cDC1 clusters or in cluster 14 CD8^+^ T vs. other CD8^+^ T cell clusters. **g-h,** The indicated CD8^+^ T cell subsets were sorted from the spleens of 3-week-old mice and stimulated with anti-CD3/anti-CD28 followed by staining for CD107a (**g**) or with PMA/ionomycin and stained for Granzyme B (**h**). **i,** CD49d^+^ CD8^+^ T cells were sorted from the spleens of 3-week-old *Clec9a^+/cre^Stat1^fl/fl^*and *Stat1^fl/fl^* mice and stimulated with PMA/ionomycin as above. The frequency of Granzyme B expression is shown. Each dot represents one mouse from 2 independent experiments, horizontal bars represent mean, error bars represent SD. Statistical analysis was performed using one-way ANOVA with Tukey’s multiple comparisons, *p < 0.05, **p < 0.01 ***p< 0.001 ****p < 0.0001.

### Chow diet boosts CXCL9 production from splenic cDC1 and the effector differentiation of food antigen specific CD8^+^ T cells

In infants increased dietary complexity correlates to systemic immune alterations, including higher plasma IFNγ levels^73,74^. To directly assess if weaning associated dietary changes influence CXCL9 production from splenic cDC1, we first induced a delay in weaning by restricting access to chow and prolonging milk diet after pups were separated from the mothers^4^ (Fig. 8a). Indeed, one week after separation from the mothers, cDC1 from mice weaned onto milk showed lower CXCL9 production than cDC1 from mice weaned normally onto chow (Fig. 8a). Similarly, IFNγ production in CD8^+^ T, NK and NKT cells was lower in mice weaned onto milk than in mice weaned onto chow (Fig. 8b). Chow contains heterogeneous plant-derived fibers and other ill-defined contents, such as lipopolysaccharide (LPS)^75,76^. These are not found in breast milk or purified mouse diets and can trigger TLR-mediated immune responses independent of the microbiota^75,76^. Indeed, CXCL9 production from spleen cDC1 and IFNγ production from CD8^+^ T, NK and NKT cells was lower in 3-week-old GF *Myd88^-/-^Trif^LPS^*^2*/LPS2*^ compared to wild type mice (Fig. 8c, d, Supp. Fig. 9a). These data suggest that during weaning dietary components in chow trigger a *Myd88/Trif*-dependent immune response independent of the microbiota, that via IFNγ production from lymphocytes is relayed to splenic cDC1 and boosts their CXCL9 production.

**Figure 8:**
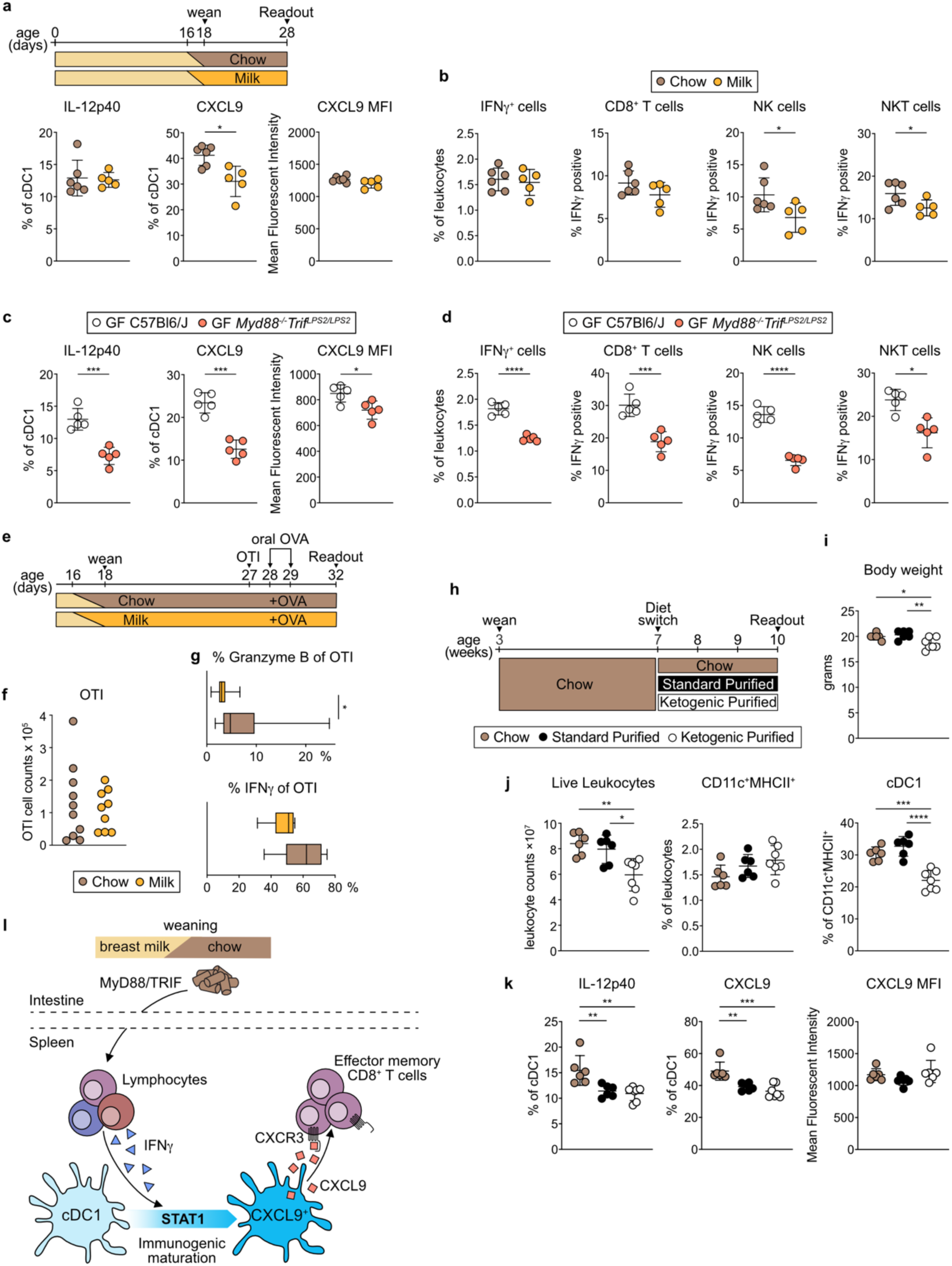
Chow diet boosts CXCL9 production from splenic cDC1 and the effector differentiation of food antigen specific CD8^+^ T cells. **a-b,** SPF mice were either conventionally weaned on day 18 or weaning was delayed by placing mice on formula milk from day 16 after birth while restricting access to chow (milk group). Exclusive feeding on formula milk was sustained until analysis on day 28. **a,** Quantification of IL-12p40^+^ and CXCL9^+^ cDC1 from indicated groups. **b,** Frequency of IFNγ production in total live leukocytes, CD8^+^ T cells, NK and NKT cells from indicated groups. (Data are representative of two independent experiments). **c-d,** Splenocytes from 3-week-old GF *Myd88^-/-^Trif^LPS2/LPS2^* or age-matched GF wild type controls were analyzed. c, Frequency of IL-12p40^+^ and CXCL9^+^ cDC1 **d**, Frequency of IFNγ^+^ leukocytes, CD8^+^ T cells, NK and NKT cells is shown. **e-g,** Weaning was delayed in SPF mice as described above. At 27 days old mice received 1×10^5^ naïve OTI cells. On the next two consecutive days, mice were gavaged with OVA and three days after the last gavage OTI T cells in spleens were analysed. **e**, Experimental scheme. **f, g,** The number of OTI cells (f) and frequency of IFNγ- and Granzyme B- producing OTI T cells in spleen is shown. OTI number and GzmB data (f, g) are pooled from three independent experiments, while IFNγ data (g) are pooled from two independent experiment. **h-k,** Mice were weaned onto chow. At 6-7 weeks of age they were either continued on normal chow or switched to a standard purified or ketogenic purified diet for 3 weeks after which splenocytes were analyzed. **h,** Experimental scheme. **i,** Body weight at readout. **j,** Spleen leukocyte counts, frequency of splenic CD11c^+^MHCII^+^ cells and frequency of cDC1 within CD11c^+^MHCII^+^ cells. **k,** Quantification of IL-12p40^+^ and CXCL9^+^ cDC1 from adult mice fed with the indicated diets for 3 weeks. Data in **g**, and **h-k** are pooled from two independent experiments, data in f is pooled from three independent experiments. Each dot represents one biological replicate, horizontal bars represent mean, error bars represent SD. Statistical analysis was performed using two-tailed Welch’s *t*-test (**a-g**), or one-way ANOVA with Tukey’s multiple comparisons **(h-k)**, *p < 0.05, **p < 0.01, ***p < 0.001, ****p < 0.0001. **l**, Model: Dietary cues present in chow induce IFNγ production from lymphocytes. In spleen, lymphocyte derived IFNγ acts via STAT1 in cDC1 to drive them into a CXCL9-producing maturation state that promotes the expansion of cytotoxic CD8⁺ effector-like T cells.

Antigen-experienced CD8^+^ T_EM_ cells in steady state SPF and GF mice are thought to arise in response to environmental antigens, such as food. Indeed, we observed an increase in the percentage and number of CD49d^+^CD44^+^ CD8^+^ T cells in spleens between two and 3 weeks of age, which correlates to the onset of weaning, i.e. when pups start eating chow (Supp. Fig. 9b). To test if weaning influences the effector phenotype of food-antigen specific CD8^+^ T cell in spleens, we either weaned mice conventionally or onto a milk diet as above. 9 days after separating pups from the mothers, we transferred naïve OTI T cells into these mice, and subsequently provided OVA as food antigen or PBS as control (Fig. 8e, Supp. Fig. 9c). Three days after the last OVA feeding, we analyzed OTI T cells in spleens. We first confirmed antigen specificity of the response in both diets, observing robust recovery of OTI T cells only in spleens of mice that had received OVA, but not PBS as control (Supp. Fig. 9d, e). Similar numbers of OTI T cells were recovered when comparing milk and chow groups (Fig. 8f, Supp. Fig. 9e). However, we observed higher Granzyme B and IFNγ production in OTI T cells recovered from spleens of mice on chow compared to milk diet, indicating more robust effector differentiation (Fig. 8g).

Finally, we asked if dietary intervention in adulthood could alter the maturation of cDC1 into CXCL9 producing cells. We weaned mice conventionally onto standard chow and at 6-7 weeks of age, we either kept mice on chow or fed them with a ketogenic purified or standard purified diet for three weeks ad libitum (Fig. 8h). Both purified diets are similarly formulated and lack complex plant-derived fibers and other potentially bioactive compounds. A standard purified diet mirrors chow in macronutrient ratio while a ketogenic purified diet shares significant metabolic similarities with breastfeeding - both are high in fat, low in carbohydrates and protein^77,78^. As expected^79^, mice fed ketogenic diet exhibited lower body weight compared to mice fed standard chow or purified control diet (Fig. 8i). We also observed reduced spleen cellularity in mice fed ketogenic diet (Fig. 8j). The frequency of CD11c^+^MHCII^+^ cells within splenocytes was similar across diets but we observed a relative reduction of XCR1^+^ cDC1 specifically in mice fed ketogenic diet compared to standard purified or chow diet (Fig. 8j), suggesting that these cells are sensitive to fat metabolism. However, CXCL9 and IL-12p40 production from cDC1 were reduced in mice fed with either of the purified diets compared to standard chow diet (Fig. 8k). Taken together, we show that IFNγ produced from lymphocytes in response to dietary components present in chow drives cDC1 into an immunogenic maturation state that promotes the expansion of cytotoxic CD8⁺ effector-like T cells during weaning (Fig. 8l).

## DISCUSSION

While the regulatory circuits that promote T cell tolerance in early life are well studied, the signals that determine immune stimulation remain elusive. Here, we identify a diet-driven, IFNγ-mediated regulatory circuit that relays dietary information to cDC1 in spleen and enables them to propagate cytotoxic effector-like CD8^+^ T cells independent of the microbiota. During weaning this circuit regulates food-antigen specific CD8^+^ T cells in a feed-forward fashion, potentially preparing the organism for new immune challenges associated with increased dietary complexity at the time of nutritional independence. cDC1 maturation remained responsive to dietary intervention in adult mice, highlighting the potential to harness dietary intervention to modulate cDC function therapeutically or in vaccination beyond the weaning period.

We identified steady state IFNγ−mediated STAT1 activation as a driving factor for a specific CXCL9^+^ maturation state of cDC1 transcriptionally distinct from CCR7^+^ cDC1. Although IFNγ signaling regulates transcriptional targets beyond *Cxcl9,* this chemokine serves as a representative marker allowing detection by flow cytometry. This cDC1 state expresses genes associated with immunogenic DC maturation after PAMP stimulation^24^ but it arises in the absence of pathogens and even under GF conditions, showing it is the result of homeostatic maturation. cDC intrinsic loss of *Stat1* reduces cytotoxic CD8^+^ T_EM_ cells in spleen, suggesting that the homeostatic maturation of cDC1 is not exclusively tolerogenic. Prior work had identified a similar cDC1 state by scRNA-seq in adult mouse spleen and trajectory analysis suggested these cells represent a transient “early mature” stage of tolerogenic cDC1 maturation into CCR7^+^ cells^25^. Whether STAT1-regulated *Cxcl9^+^* cDC1 are part of a single linear maturation trajectory or represent a distinct maturation state of cDC1 remains to be determined. CD8 T cell responses are subject to exact spatiotemporal regulation involving IFNγ and CXCL9 cross-talk also in infections and tumours^80–85^. Tumor-resident CXCL9-expressing cDC1 that lack CCR7 induce the local activation of protective anti-cancer CD8^+^ T cell responses^17,86^. Similarly, our i.v. labeling and transcriptional data indicate that the interaction of CD8^+^ T cells and homeostatically matured CXCL9^+^ cDC1 takes place outside the T cell zone, likely in blood exposed regions, such as the marginal zone or red pulp, known sites for the regulation and bystander activation of memory CD8^+^ T cells^80–83^. This localization may strategically place cDC1 allowing them to acquire antigens or other signals from blood that further promote their interaction with CD8^+^ T cells.

*Stat1* deletion did not abrogate CXCL9 production in all cDC1, indicating that additional signals regulate CXCL9 expression in cDC1. Indeed, loss of STING causes a reduction of CXCL9 production from cDC1 in steady state SPF mice^25^. STING-mediated CXCL9 production could be linked to recognition of self-DNA from apoptotic cells or recognition of bacterial DNA that can be delivered into host cells from microbiota-derived membrane vesicles^25,87,88^. Both signals would trigger the cytosolic cGAS/STING pathway, which can induce production of IFN-I and subsequent CXCL9 production from cDC1^25,88^. Signals driving CXCL9 in cDC1 could differ by age and CXCL9 production in cDC1 may even be regulated in a tissue-specific manner.

Dietary supplementation and nutritional interventions are increasingly used to modulate immunity to reduce inflammation or alter immune responses, yet nutrition remains a poorly understood regulator of immunity, in part because altering diet also changes host-microbial cross talk^89–92^. Chow contains various micronutrients and other poorly defined bioactive components, including traces of microbial components, such as LPS, or beta-glucan, a soluble fiber found in the cell walls of grains^93^ that can trigger PRR signaling and cytokine production from immune cells^76,94–96^. Dietary LPS for instance activates mucosal immunity and shapes the intestinal IgA repertoire^75^. Our data strongly suggest that Myd88-TRIF activating bioactive dietary components initiate the here identified IFNγ-mediated regulatory circuit, although we cannot exclude that mechanical forces related to dietary consistency trigger Myd88-TRIF dependent immune activation. Further work is needed to define the exact dietary components sensed, if they are sensed locally in the intestine or elsewhere, and the exact cell types involved. Vitamin C has been shown to directly modify lysine residues in STAT1, which prolongs STAT1 phosphorylation and enhances anti-tumor immunity in mice^97^. STAT1 induces IFNγ production^51^, raising the possibility that dietary supplementation of Vitamin C could be used to promote the immunogenic maturation of cDC1 by inducing IFNγ production from lymphocytes or by promoting STAT1 phosphorylation in cDC1 themselves. Mice can synthesize Vitamin C from glucose, making this pathway irrelevant in the murine model, however, humans require Vitamin C from food and dietary supplementation with Vitamin C is commonly used to boost anti-viral immunity^98^. If a specific IFNγ producing lymphocyte population drives CXCL9 production from cDC1 remains to be determined, as well as whether these IFNγ-producing lymphocytes, specifically CD8^+^ T cells, originate from the intestine. Since diet can modulate the CXCL9^+^ cDC1 state in adult mice, it will be interesting to determine if diet-induced changes in IFNγ production regulate cDC1 even after periods of malnutrition or if chow diet is introduced at later stages in life.

Steady state CD49d^+^CD8^+^ T_EM_ cells likely recognize environmental or food-derived antigens^99,100^. At three weeks of age, these cells showed transcriptional and functional similarity with cytotoxic CD8^+^ T_EM_ cells, including ability to rapidly degranulate and high expression of Granzymes and *Cx3cr1*^71^. The few CD8^+^ T_EM_ cells that are found in spleens of *Clec9a^+/cre^Stat1^fl/fl^* mice still produce Granzyme B upon stimulation, suggesting intact priming. We believe that these cells are not primed in the spleen, but rather are attracted by IFNγ-stimulated CXCL9⁺ cDC1 to splenic niches after being initially primed elsewhere. Antigens encountered via the diet normally induce T cell tolerance, although such tolerance mechanisms can be suspended depending on the context in which dietary antigens are sampled^101–103^. Our data suggest that the immune system systemically relays dietary cues during weaning, allowing it to tailor the T cell response against environmental antigens to the need of the developing organism, striking a balance between effector immunity and T cell tolerance. This is supported by the observation that food-antigen specific CD8^+^ T cells were more cytotoxic in spleens of weanling mice on chow compared to milk diet. Effective desensitization against food allergens by oral immunotherapy has been linked to an increased frequency of cytotoxic CD8^+^ T_EM_ cells, specifically in peripheral blood of patients with sustained unresponsiveness over those with recurring allergy^104^. Our data suggest that diet may influence the effectiveness of oral immunotherapy by first influencing T cell priming in the gastrointestinal tract and subsequently forwarding information to distant sites to shaping the functions of antigen presenting cells in a feed forward manner.

Together, our study identifies a previously unrecognized, microbiota-independent regulatory circuit by which dietary cues are relayed through IFNγ to imprint spleen-resident cDC1 with an immunogenic maturation program. This circuit enables cDC1 to shape cytotoxic effector memory CD8⁺ T cells in early life to tailor the T cell pool to developmental stage. These findings redefine the landscape of steady-state cDC1 maturation, challenging the prevailing view that homeostatic cDC1 maturation primarily promotes tolerance. By uncovering a critical link between nutrition and cDC function that remains responsive to dietary interventions in adulthood, our work opens new avenues for leveraging dietary interventions to modulate cDC-driven immunity in vaccination or disease.

## MATERIALS AND METHODS

### Mice

Clec9a^tm2.1(icre)Crs^ (*Clec9a^Cre^*) (Jackson Laboratory Stock No: 025523), Gt(ROSA)26Sor^tm9(CAG-tdTomato)Hze^ (*Rosa26^lox-STOP-lox-tdtomato^*) (Jackson Laboratory Stock No: 007909), Ifnar1^tm1Agt^ (*Ifnar^−/−^)* (MMRRC Stock No: 32045-JAX), B6(Cg)-Ifnar1tm^1(flox)Uka^ (*Ifnar1^fl/fl^*), Tg(Itgax-cre)1-1Reiz (*Itgax^cre^*) (Jackson Laboratory Stock No: 018967), Stat1^tm1.1Mmul^ (*Stat1^fl/fl^*), C57BL/6-Tg(Nr4a1-EGFP/cre)820Khog-Ptprca (*Nr4a1^eGFP^*) (Jackson Laboratory Stock No: 016617), OTI mice (C57BL/6-Tg(TcraTcrb)1100Mjb/J, Jackson Laboratory stock no: 003831) and C57BL/6JRccHsd were bred and maintained at the Biomedical Center, LMU Munich. B6.129S7-Ifngr1^tm1Agt^/J mice were maintained at the VIB Center for Inflammation Research Ghent (Jackson Laboratory Stock No: 003288). C57BL/6J wildlings were created through inverse germ-free rederivation as described^57^. SPF mice were maintained in individually vented cages with a 12 h dark/light cycle. Wildling mice were bred at the Medical Center, University of Freiburg, Germany. Germ-free animals were bred at the ZIEL institute for Food & Health TUM, Freising, Germany in isolators and germ-free status was routinely monitored. Germ-free *Myd88^-/-^Trif^LPS2/LPS^*^2105^ mice were bred and maintained in flexible-film isolators at the Clean Mouse Facility, University of Bern, Switzerland. Food and water were provided ad libitum. Unless otherwise specified mice were housed on chow as a grain-based diet. Chow diets may exhibit considerable batch-to-batch variation but are used in most experimental facilities worldwide so that we did not specifically normalize chow diets across facilities. Both male and female mice were used in this study. Littermates were used in experiments unless otherwise stated. All animal procedures were performed in accordance with national and institutional guidelines for animal welfare and approved by the respective authorities.

### Delayed weaning

We delayed weaning following an established protocol^4^. Litter sizes between the formula-fed (milk) and chow-fed groups were equalized one week after birth. Between postnatal days 16 and 18, solid food (chow) was removed from the cage of the milk-fed group and the pups received 15µL of Optima Pet Milk (TVM) by oral gavage 8 times per day. The dam was also fed with formula milk provided in sterile bottles during this time. The chow group had continuous access to standard chow diet and served as the control. 18 days after birth the dams were removed from both cages and the pups in the formula-fed group continued to receive formula milk only via bottle. Bottles were replaced three times a day to prevent bacterial contamination, cages were changed twice a day to control for coprophagy. Animals from both groups were analyzed at 4 weeks of age.

### Food antigen response of OTI T cells

Adoptive transfer of OTI T cells and oral OVA administration was performed within the delayed-weaning experimental setting described above. 9 days after being separated from the mothers, CD45.2 congenic pups were injected intravenously with 1×10⁵ naïve OTI T cells isolated from adult CD45.1/2 congenic OTI mice using the MojoSort™ Mouse CD8 Naïve T Cell Isolation Kit (BioLegend). 1 and 2 days after adoptive OTI T cell transfer mice were gavaged with 50 mg OVA (Grade III, Sigma, A5378) in 50 µL PBS or with 50 µL PBS as control. 3 days after final OVA administration mice were analysed. A CD45.2⁺ mouse that did not receive OTI T cells served as a control to set the gate for injected OTI T cells.

### Ketogenic diet

6–7-week-old female mice were fed standard chow diet (Ssniff), standard purified diet (AIN-93M, powder, Bio-Serv) or ketogenic purified diet (AIN-76A Modified, High Fat, Paste, Bio-Serv) for 3 weeks ad libitum (Supp. Table 7). All diets used in this study were sterilized either by autoclaving or irradiation.

### Cell isolation

Mice were sacrificed by cervical dislocation. Mice younger than 3 weeks were sacrificed by decapitation. Spleens were isolated, minced and enzymatically digested in 1mL RPMI containing 200U/mL Collagenase IV (Worthington) and 0.2mg/mL DNaseI (Roche) for 30min at 37°C while shaking (180 rpm), then passed through a 70μm strainer. The tubes were then filled with ice cold FACS buffer (PBS 1% fetal calf serum (FCS), 2.5mM EDTA, with 0.02% sodium azide) and centrifuged. Subsequently, erythrocytes were osmotically lysed in Ammonium-chloride-potassium (ACK) buffer (MilliQ water with 1mM EDTA, 1.55M NH_4_Cl and 100mM KHCO_3_) for two minutes at 4°C, and washed with FACS buffer. Cell pellets were resuspended in FACS buffer and passed again through a 70μm strainer before further analysis. For functional analyses and scRNA-seq experiments PBS containing 1% FCS and 2.5mM EDTA was used for cell isolation.

### Flow Cytometry

4×10^6^ cells were stained in 50µL FACS buffer with anti-mouse CD16/32 (Fc-block) for 10 min at 4°C. 50µL of a 2x mastermix containing antibodies against surface epitopes and fixable viability dye eFluor™ 780 (Thermo Fisher Scientific) was added and cells were incubated at 4°C for 25 min. Cells were then washed twice and resuspended in FACS buffer for analysis. After surface staining, intracellular cytokine staining was performed using the Intracellular Fixation & Permeabilization Buffer Set (Thermo Fisher Scientific); intranuclear markers and Granzyme B were stained using the Foxp3 Transcription Factor Staining Set (Thermo Fisher Scientific) according to the manufacturer’s instructions. CountBright™ Absolute Counting Beads (Thermo Fisher Scientific) were added to obtain cell counts as previously described^106^. For intracellular cytokine staining of lymphocytes, splenocytes were stimulated with 10ng/mL phorbol 12-myristate 13-acetate (PMA) and ionomycin (1µg/mL) for 5h at 37°C. Brefeldin A (5µg/mL) was added for the last 3h. Prior to IL-12p40 and CXCL9 staining splenocytes were incubated for 4h with Brefeldin A (5µg/mL) and Monensin (2µM) at 37°C. MAIT cells were stained with BV421-conjugated mouse MR1-5-OP-RU tetramers at room temperature for 30 min. 6-FP-loaded MR1 tetramers were used as a negative control^107^. Antibodies used for flow cytometry are provided in Supp. Table 5. Data was collected on a LSR Fortessa (BD Biosciences) using BD FACSDiva Software (BD BioSciences, version 8) and analyzed using FlowJo software (Tree Star, Inc.). Mean fluorescence intensity was calculated as the geometric mean of the indicated fluorescent parameter. Cell sorting was performed on a FACSAria Fusion (BD Biosciences).

### In vitro stimulation with PAMPs

For in vitro stimulation of cDC1 with PAMPs, splenocytes from spleens of 2-week-old or adult mice were enriched using CD11c MicroBeads (Miltenyi) according to the manufacturer’s recommendation. 2-4 spleens from young mice of the same sex were pooled to obtain sufficient cell numbers. Cells were then stained as above in PBS containing 1%FCS and 2.5mM EDTA and cDC1 were sorted into the PBS containing 10% FCS according to the gating strategy in Supp. Fig. 1a). 35,000 XCR1^+^ cDC1 were stimulated with 0.5µg/mL CpG-B ODN 1826 (Sigma), 10µg/mL Zymosan A (Sigma), 100µg/mL Zymosan Depleted (InvivoGen), or 10µg/mL poly(I:C) LMW (InvivoGen) in a total volume of 50µL RPMI containing 10% FCS, 1% Penicillin/Streptomycin, 1% non-essential amino acids, 1% sodium pyruvate, 1% L-Glutamine, 0.05mM β-mercaptoethanol. After 20 hours cells were pelleted and cytokines in supernatants were quantified using LEGENDplex™ Mouse Inflammation Panel and LEGENDplex™ Mouse Cytokine Panel 2 for IL-12p40 (both Biolegend).

### T cell degranulation assay

Splenocytes from 3-week-old mice were stained with FITC-conjugated antibodies against CD19, Ly6G, Ter119 and CD4. Splenocytes were then depleted for FITC labelled cells using anti-FITC magnetic beads (Miltenyi) and LS columns (Miltenyi) according to manufacturer’s instructions. Subsequently, cells were washed and surface stained for surface epitopes as described above and then sorted into PBS containing 10% FCS. 9.000 naïve CD8^+^ T cells (CD62L^+^CD44^neg^), CD44^+^CD49d^+^ and CD44^+^CD49d^neg^ CD8^+^ T cells were sort-purified and incubated in 96-well v-bottom plates coated with 1µg/ml LEAF-purified anti-CD3e (Biolegend) at 4°C overnight, in the presence of soluble anti-CD28 (1µg/ml) for a total of 5h at 37°C. Anti-CD107a (Biolegend) was added to the culture along with Brefeldin A and Monensin for the last three hours of culture. The cells were then re-stained with viability dye. For granzyme B staining, 9.000 sorted cells were incubated with PMA and ionomycin for a total of 5h. Brefeldin A was added for the last three hours.

### Quantitative Real-Time PCR

RNA was isolated from spleens using RNeasy Midi Kit (Qiagen) and subsequently treated with DNase I to remove any residual genomic DNA, followed by enzymatic inactivation using the TURBO DNA-free™ Kit (Invitrogen). Complementary DNA (cDNA) was synthesized using Superscript III reverse transcriptase (Invitrogen) according to manufacturer’s instructions. 2 µg RNA was used for reverse transcriptase reaction. In parallel a “no RT control” reaction was performed with the same 2 µg RNA input but no-reverse transcriptase was added. Quantitative Real time PCR performed using SYBR Green Fast master mix (Invitrogen) according to the manufacturer’s instructions, on a Real-Time PCR system (Applied Biosystems) using primers listed in supplementary table 6. For primers that are not exon spanning qRT-PCR was performed on cDNA samples and respective no-RT controls. Only samples showing specific amplification in the cDNA reaction with no detectable product in the no-RT control were quantified. Quantification was performed by relative standard curve method and target gene expression was normalized to *Actb*. For visualization purposes a normalization factor (x1000) was applied to the relative expression values.

### Single cell RNA sequencing

Splenocytes from 1-week (n=5), 3-week (n=4), 4-week (n=3) and 6-week (n=3) old *Clec9a^+/cre^Rosa^TOM^* mice were isolated and stained with FITC-conjugated antibodies against CD19, CD3e and Ter119. Cells were then depleted of FITC-labelled cells using anti-FITC magnetic beads (Miltenyi) and LS columns (Miltenyi) according to manufacturer’s instructions. Depleted samples were labelled with barcoded antibodies (anti-MHCI, anti-CD45, TotalSeq™-B0305/B0310 anti-mouse Hashtag Antibody, Biolegend) according to time point, and CD11c^+^MHCII^+^ cells were sorted into PBS containing 10% FCS (Supp. Fig. 2a). Sorted CD11c^+^MHCII^+^ cells from two time points, labelled with different barcoded antibodies, were pooled, pelleted, and then resuspended to 700 cells/µL in PBS containing 0.04% bovine serum albumin (BSA). Each pool was then loaded onto a separate reaction of a Chromium chip (10X Genomics). Gene expression and cell surface protein libraries were prepared using the Chromium Next GEM Single Cell 3’ Reagent kit (10X Genomics). Concentration and purity of the libraries was assessed with a TapeStation (Agilent). Libraries were multiplexed and sequenced using the recommended sequencing depth on a NextSeq1000 (Illumina).

For scRNA-seq of splenic CD11c^+^MHCII^+^ DCs and CD90^+^ cells from 3-week-old *Clec9a^+/cre^Stat1^fl/fl^*(n=2) and *Stat1^fl/fl^* littermates (n=2) littermates, splenocytes were isolated and depleted of CD19, Ly6G and Ter119 positive cells as above using magnetic beads. Depleted samples were then labelled with barcoded antibodies according to genotype, as described above. After sorting, purified DCs from both genotypes were pooled and resuspended to 1000 cells/µL in PBS containing 0.04% BSA. In the same manner, CD90^+^ cells from both genotypes were pooled and resuspended. Each of the two pools was then loaded onto a separate reaction well of the Chromium Chip (10X Genomics). Library generation, quality control and sequencing were performed as described above.

### scRNAseq analysis

Sequencing data were processed using 10X Genomics Cell Ranger v6.0.0 pipeline and mapped to the mouse genome (mm10) customized to include the sequence of iCRE (GenBank ID: AY056050.1), Tomato (GenBank ID: AY678269.1) and the predicted transcript of the unrecombined Rosa locus. For the dataset across age, raw count matrices and cell surface protein information were aggregated with the Cell Ranger aggregate pipeline. The resulting gene-barcode matrix was loaded into R (v4.3.2) using the *Seurat* (v4.3.0) package. Genes detected in <3 cells and cells expressing <1500 genes or >5% mitochondrial genes were excluded from further analysis. Cells were scored based on the expression of cell cycle associated genes and cell cycle regression was performed according to Seurat’s “Cell-Cycle Scoring and Regression” vignette. Cells of different time points were identified by cell surface protein information of barcoded antibodies. Doublets were identified by dual labelling with both barcoded antibodies and also excluded, along with cells not exhibiting sufficient barcode labelling levels to allow for a clear distinction. The *sctransform* package was used to normalize, scale and find variable features of the dataset. Dimensional reduction by UMAP was based on the first 60 principal components. Louvain clustering was used to cluster the data in an unsupervised manner. The resolution was chosen such that RORγt^+^ DC clustered separately. Differentially expressed genes between clusters were identified using the FindAllMarkers and FindMarkers commands of the *Seurat* package. *AUCell* (v.3.18) package was used to score cells for enrichment of published gene signatures (Supp. Table 1). Gene set enrichment analysis of hallmark gene sets was performed using the *singleseqgset* package.

For the dataset containing CD11c^+^MHCII^+^ cDCs and CD90^+^ cells from *Clec9a^+/cre^Stat1^fl/fl^*mice and *Stat1^fl/fl^* littermates, DC and lymphocyte libraries were analyzed separately with *Seurat* in R. The *SoupX* package was used to decontaminate cells from ambient RNA and the *scDblFinder* package was used to identify and exclude putative doublets. HTO based doublet exclusion was performed as described above. For the cDC library, genes detected in <3 cells and cells expressing <1000 genes or >5% mitochondrial genes were excluded from further analysis. For the CD90^+^ lymphocyte library, genes detected in <3 cells and cells expressing <1000 genes or >10% mitochondrial genes were excluded from further analysis. Louvain clustering was used to cluster both, DC and CD90^+^ lymphocyte datasets, in an unsupervised manner. Identification of differentially expressed genes, scoring of published gene signatures and GSEA was performed as described above. Interactome analysis was performed using *Community*^70^ using aggregated gene expression data of DC and T cell clusters.

### Multiome computational analysis

Cells annotated as cDC1 in the multiomic dataset of splenic CD11c^+^MHCII^+^ cells and MHCII^+^ ILC3s^40^ were computationally isolated, followed by dimensional reduction and unsupervised clustering according to the standard scanpy work flow (https://scanpy-tutorials.readthedocs.io/en/latest/pbmc3k.html). Gene set enrichment analysis between cDC1 from 2-week-old and adult mice was performed using *gseapy* package with default parameters^108^.

**Intravenous labelling** was performed as described previously^25,109^. 3μg Pacific Blue-conjugated anti-CD45.2 was injected intravenously into 3-week-old mice. 2 minutes post injection, the mice were sacrificed by cervical dislocation, and flow cytometry of splenocytes was performed.

### Statistical analysis

Statistical analyses were performed in Prism 10 software (GraphPad) using two-tailed t-test with Welch’s correction (unless otherwise stated). Percentage data were logit-transformed prior to statistical testing. For multiple comparisons one-way analysis of variance with Tukey’s test (unless otherwise stated) was performed. For comparing expression levels and AUCscores in scRNA-seq datasets statistical analysis was performed using multiple t-tests corrected for multiple comparisons using the Holm-Šídák method. For analyses of food antigen specific T cells linear mixed-effects models (R/lme4 version 2.0-1) were used to account for batch effects, with experiment included as a random intercept and condition as a fixed effect. This approach estimates the contribution of inter-experimental variability and tests condition effects within experiments rather than across pooled data.

## ACKNOWLEDGEMENTS

We thank Anne Krug and members of the Schraml lab for helpful discussions and critical reading of the manuscript. We also thank Amina Sayed for technical help. We acknowledge the Core Facilities for Flow Cytometry, Bioimaging and Animal Models at the Biomedical Center, LMU Munich for providing equipment and expertise. High-throughput sequencing was performed by the Laboratory for Functional GenomeAnalysis (LAFUGA) of the LMU Munich. This work has been funded by an ERC Starting Grant (ERC-2016-STG-715182) and by the Deutsche Forschungsgemeinschaft (DFG, German Research Foundation) - TRR 359—Project number 491676693 (projects A07, B05 and B09, FOR2599 (Project P03, SCHR 1444/2-1). D.A. was supported by a YLSY Doctoral Scholarship from the Republic of Türkiye Ministry of National Education. J.P.B. acknowledges support by the DFG (project numbers 461704785 – SPP2306, 424926990, 442405234 and 449174900) and the Wilhem Sander-Stiftung (project number 2024.133.1).

## AUTHOR CONTRIBUTION

R.S., D.A., K.R.R., H.N., N.E.P., D.M., J.V., S.N., M.P.R, and N.N. performed experiments and generated data. R.S. and M.L.R. performed bioinformatic analyses. D.H, S.S. provided germ-free mice, A.K. and D.S. provided *Ifnar1* global and conditional knock out mice, M.P. and A.G. provided wildling mice, S.J. and J.V. provided *Ifngr1^-/-^* mice, S.C.G.-V. provided germ-free *Myd88^-/-^Trif^LPS2/LPS2^* mice. C.S., A.K., M.C.T., T.S., C.S., M.P., K.B., and J.P.B contributed new reagents/analytic tools/scientific input. R.S., D.A., and B.U.S. wrote the paper. B.U.S. conceptualized and supervised the study.

## DATA AND MATERIALS AVAILABILITY

Single-cell RNA sequencing data have been deposited in BioStudies (S-BSST1933, S-BSST1937). All study data are included in the article and/or supporting information.

