## Supplemental information for "Periweaning diet-induced activation of an IFNγ-mediated regulatory circuit promotes the homeostasis of cytotoxic CD8^+^ T Cells"

### Supplemental Figure 1:

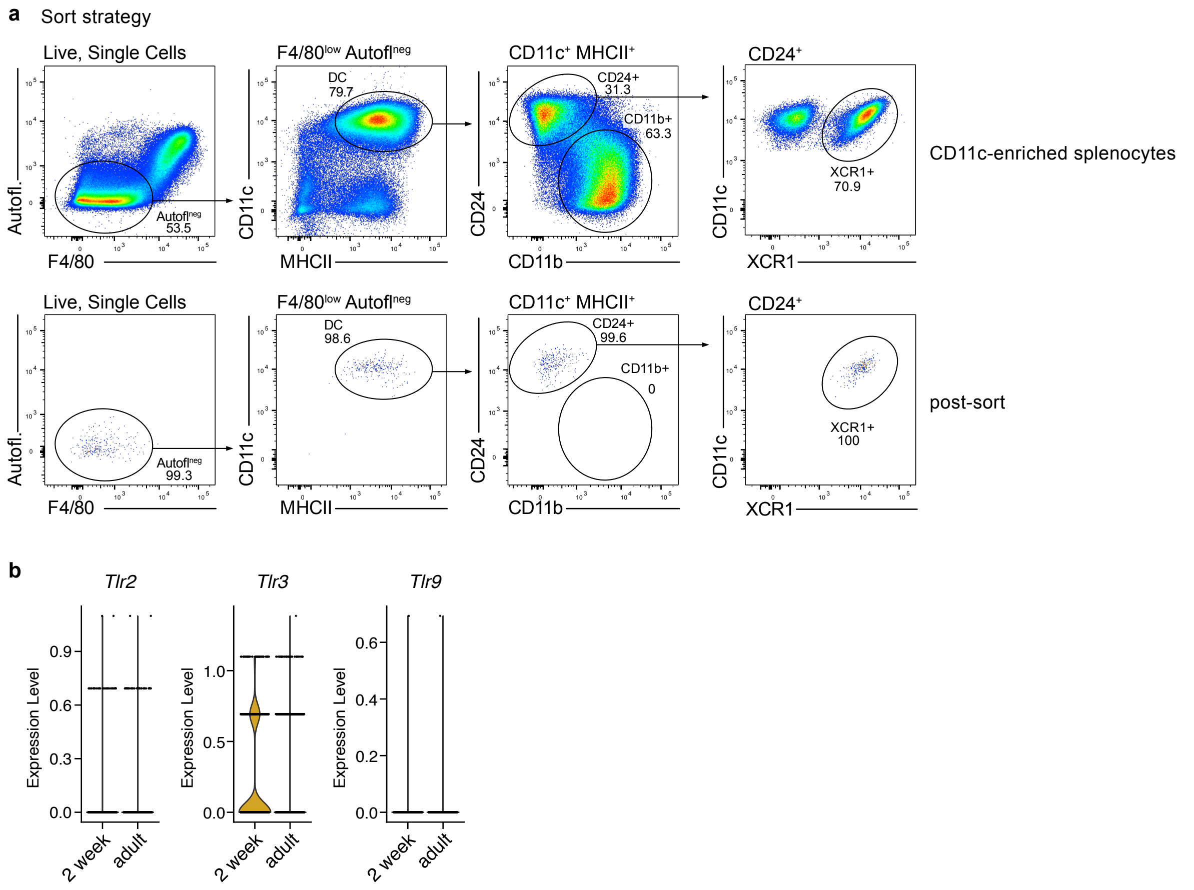

**Supplemental Figure 1: Representative sort strategy and purity of splenic cDC1**

**a,** Representative gating strategy for $XCR1^+$ cDC1 sorted from CD11c-enriched splenocytes of adult mice. Live leukocytes were gated after exclusion of doublets and red pulp macrophages were excluded as autofluorescent $F4/80^{high}$ cells. Then, $CD11c^+MHCII^+$ cells were gated and cDC1 identified as $CD24^+CD11b^{neg}XCR1^+$ cells. Top panels show sample before sorting, bottom panels show post-sort purity. **b**, Expression levels of *Tlr2*, *Tlr3* and *Tlr9* in cDC1 from 2-week-old and adult mice as determined in the scRNA/ATAC seq data from Fig. 1e-f.

### Supplemental Figure 2:

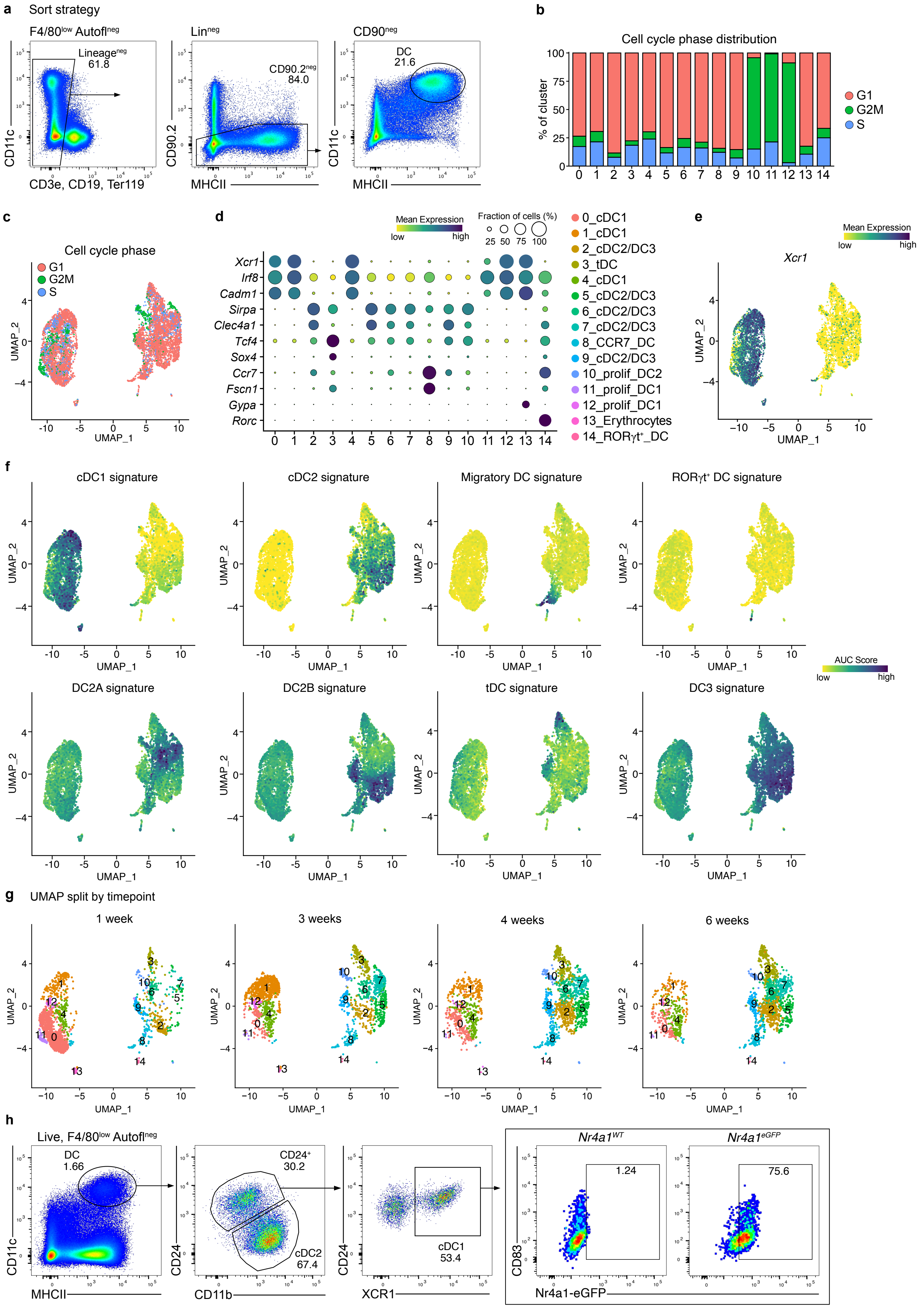

**Supplemental Figure 2: scRNA-seq analysis of dendritic cells across age**

**a-g,** CD11c$^{+}$MHCII$^{+}$ cells were sorted from spleens of mice at 1, 3, 4 and 6 weeks of age for scRNA-seq. **a,** Representative sort strategy showing cells from 6-week-old mice. CD19, CD3e and Ter119 cells were depleted using anti-FITC beads. From remaining live leukocytes F4/80$^{low}$autofluorescence$^{neg}$ cells were gated and remaining FITC$^{+}$ cells excluded. Then CD90.2$^{neg}$ cells were gated and CD11c$^{+}$MHCII$^{+}$ cells were sorted. **b, c,** Distribution of cell cycle phase across clusters from UMAP in Fig. 2b. **d,** Bubble plot of selected marker genes used for cell type identification. **e,** Expression levels of *Xcr1* on the UMAP display. **f,** UMAP display of AUC scores for the indicated gene signatures used for cluster identification. **g,** UMAP display projecting cells for the individual time points separately. **h,** Representative gating strategy for the identification of cDC1 in spleen. Live, autofluorescent negative cells from 7-week-old *Nr4a1*$^{eGFP}$ and WT mice were gated, followed by identification of CD24$^{+}$XCR1$^{+}$ cells as shown. GFP-positive cDC1 were identified based on a wildtype control.

### Supplemental Figure 3:

**a** 2 weeks spleen

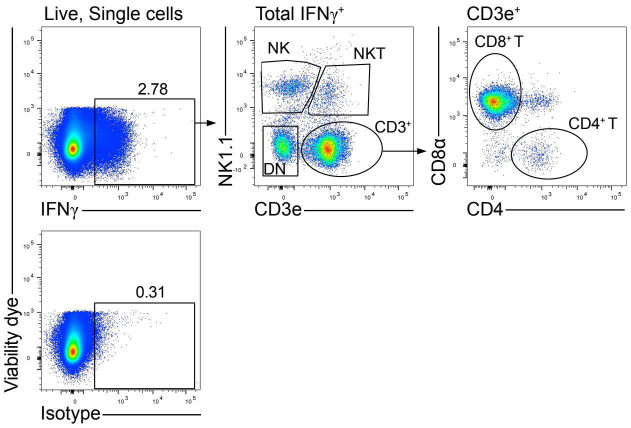

**Supplemental Figure 3: Flow cytometric analysis of IFNγ production by splenic immune cells**

**a,** Splenocytes from mice of difference ages were treated with PMA/Ionomycin for 5 hours, Brefeldin A was added for the last 3 hours. Then intracellular cytokine staining was performed. Within live leukocytes, IFNγ$^{+}$ cells were gated based on an isotype-matched control antibody (bottom panel). Within IFNγ$^{+}$ cells, NK cells (NK1.1$^{+}$CD3e$^{neg}$), NKT cells (NK1.1$^{+}$CD3e$^{+}$), T cells (NK1.1$^{neg}$CD3e$^{+}$) and NK1.1$^{neg}$CD3e$^{neg}$ cells (DN: double negative cells) were identified. T cells were further divided into CD8$^{+}$ and CD4$^{+}$ T cells. Representative gating strategy from 2-week-old mice is shown.

### Supplemental Figure 4:

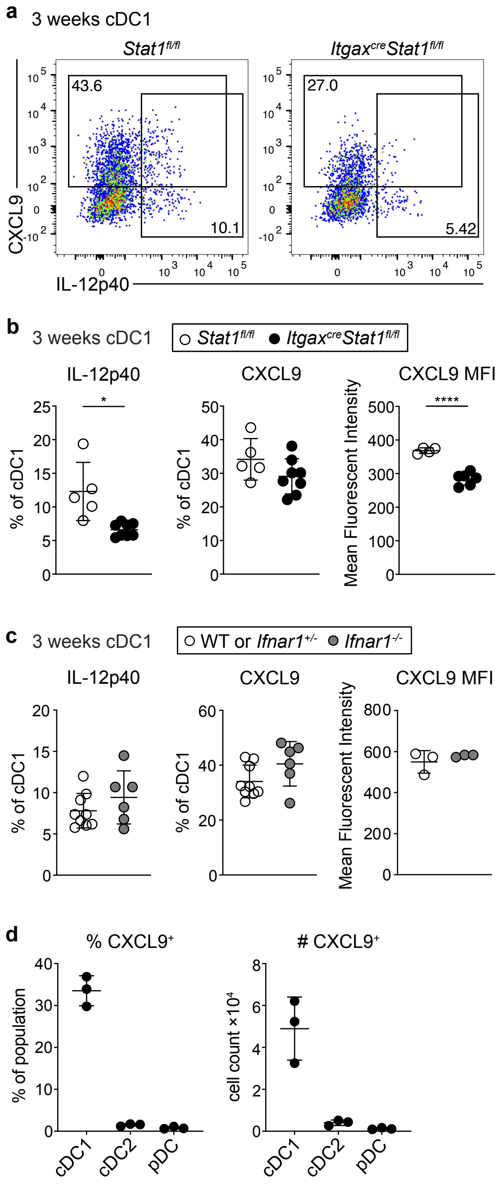

**Supplemental Figure 4: CXCL9-production from cDC1 is *Ifnar1*-independent and ketogenic diet induces changes to body weight and spleen cellularity**

**a-d** Splenocytes from 3-week-old mice of the indicated genotypes were cultured for 4 hours with Brefeldin A and Monensin. $XCR1^{+}$ cDC1 were then analyzed for cytokine production by flow cytometry. **a,** Representative gating. **b,** Quantification of $IL\text{-}12p40^{+}$ and $CXCL9^{+}$ cDC1 from 3-week-old $Itgax^{cre}Stat1^{fl/fl}$ and $Stat1^{fl/fl}$ littermate controls. **c**, Quantification of $IL\text{-}12p40^{+}$ and $CXCL9^{+}$ cDC1 from 3-week-old and $Ifnar1^{+/-}$ or $Ifnar1^{-/-}$ mice. **d**, CXCL9 production in cDC1, cDC2 and pDC from 3-week-old wildtype mice after 4 hours of culture with Brefeldin A and Monensin. Each dot represents one mouse, horizontal bars represent mean, error bars represent SD. Statistical analysis was performed using two-tailed Welch's *t*-test, $*p < 0.05$, $**p < 0.01$, $***p < 0.001$, $****p < 0.0001$.

### Supplemental Figure 5:

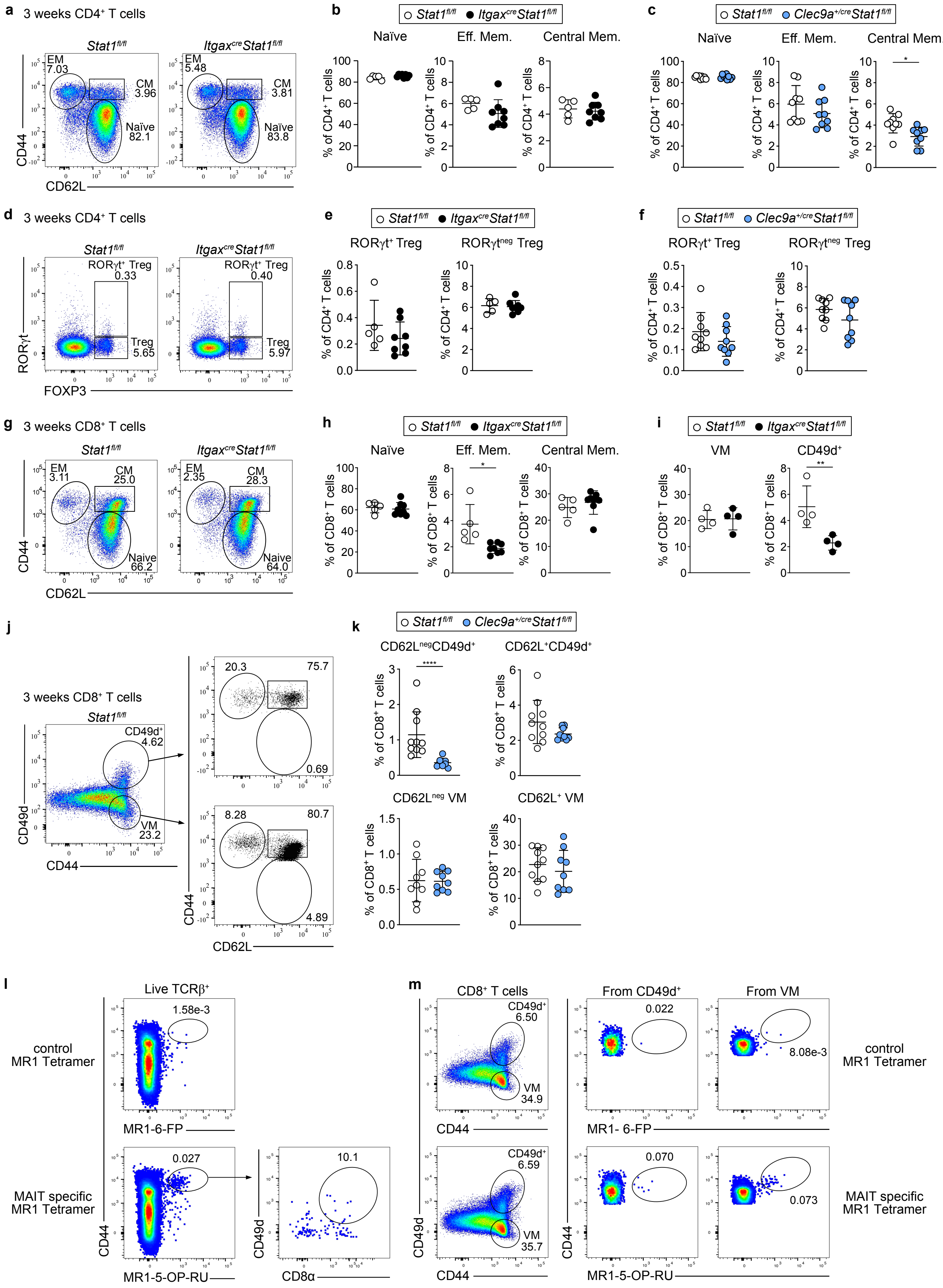

**Supplemental Figure 5: Flow cytometry of spleen T cells in 3-week-old mice**

**a-c,** $CD4^+$ T cells in the spleen of 3-week-old *Itgax*$^{cre}$*Stat1*$^{fl/fl}$ and *Stat1*$^{fl/fl}$ littermate controls and *Clec9a*$^{+/cre}$*Stat1*$^{fl/fl}$ and *Stat1*$^{fl/fl}$ were divided into effector memory ($CD44^+CD62L^{neg}$), central memory ($CD44^+CD62L^+$) and naïve ($CD44^{neg}CD62L^+$) populations. **a,** Representative gating strategy. **b-c,** quantification of the indicated populations in *Itgax*$^{cre}$*Stat1*$^{fl/fl}$ and *Stat1*$^{fl/fl}$ littermate controls **(b)** and *Clec9a*$^{+/cre}$*Stat1*$^{fl/fl}$ and *Stat1*$^{fl/fl}$ littermate controls **(c)**. **d-f,** From total $CD4^+$ T cells, $Foxp3^+$ and $Foxp3^+ROR\gamma t^+$ Tregs were gated and quantified. **d,** Representative gating strategy. **e,** Quantification of $CD4^+$ Treg populations in 3-week-old *Itgax*$^{cre}$*Stat1*$^{fl/fl}$ and *Stat1*$^{fl/fl}$ littermate controls **f,** Quantification of $CD4^+$ Treg populations in 3-week-old *Clec9a*$^{cre}$*Stat1*$^{fl/fl}$ and *Stat1*$^{fl/fl}$ littermate controls. **g-h,** $CD8^+$ T cells in the spleen were divided into effector memory ($CD44^+CD62L^{neg}$), central memory ($CD44^+CD62L^+$) and naïve ($CD44^{neg}CD62L^+$) populations. Representative gating (**g**) and quantification (**h**) of $CD8^+$ T cell subsets in spleens of 3-week-old *Itgax*$^{cre}$*Stat1*$^{fl/fl}$ and *Stat1*$^{fl/fl}$ littermate controls. **i**, Quantification of splenic $CD49d^+$ and VM CD8+ T cell populations in 3-week-old *Itgax*$^{cre}$*Stat1*$^{fl/fl}$ mice and *Stat1*$^{fl/fl}$ littermate controls. **j-k,** Splenocytes from 3-week-old *Clec9a*$^{+/cre}$*Stat1*$^{fl/fl}$ and *Stat1*$^{fl/fl}$ mice were isolated and $CD8^+$ T cells were gated as $CD49d^+CD44^+$ and $CD49d^{neg}CD44^+$ (VM). Within these populations cells were divided into central memory ($CD44^+CD62L^+$) and effector memory ($CD44^+CD62L^+$) cells (**j**). Subsets were quantified as percentage of total $CD8^+$ T cells (**k**). **l,** MAIT cells were identified in spleens of 3-week-old wild type mice as MR1-5-OP-$RU^+$ cells within $TCR\beta^+$ splenocytes as shown. 6-FP loaded MR1 tetramer was used as a control. $CD8\alpha^+CD49d^+$ cells within MAIT cells are shown. **m,** $CD44^+CD49d^+$ or $CD44^+CD49d^{neg}$ $CD8^+$ T cells were gated and then MR1-5-OP-RU signal is shown within these populations. Each dot represents one mouse, horizontal bars represent mean, error bars represent SD. Statistical analysis was performed using two-tailed Welch's *t*-test, $*p < 0.05$, $**p < 0.01$, $***p < 0.001$, $****p < 0.0001$.

### Supplemental Figure 6:

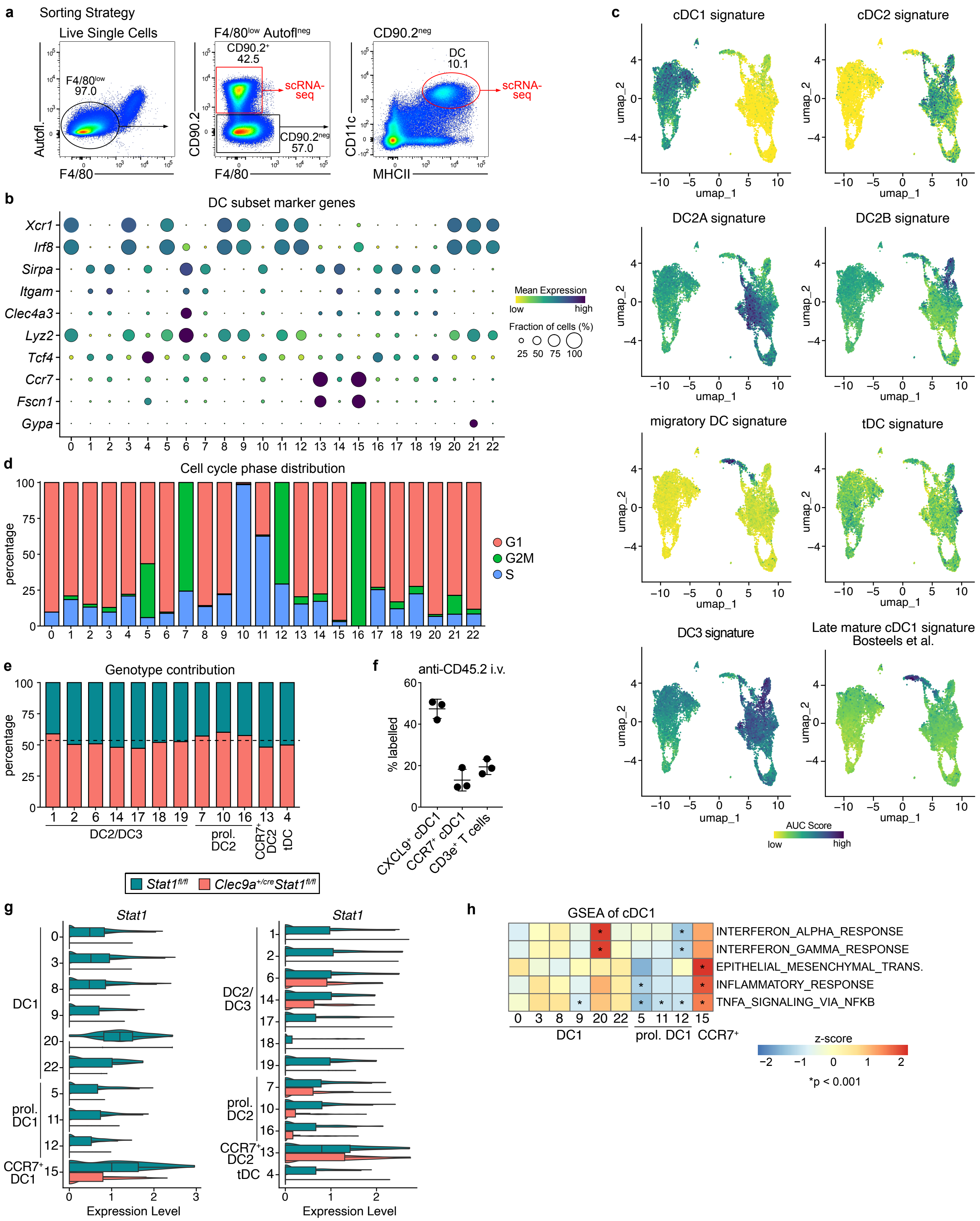

**Supplemental Figure 6: scRNA-seq of spleen cDC from 3-week-old *Clec9a$^{cre}$Stat1$^{fl/fl}$* mice**

**a,** Sort strategy for splenic CD90.2$^{+}$ cells and CD11c$^{+}$MHCII$^{+}$ cells from the spleens of 3-week-old *Clec9a$^{+/cre}$Stat1$^{fl/fl}$* and *Stat1$^{fl/fl}$* littermates. Live leukocytes were gated and red pulp macrophages excluded as F4/80$^{high}$autofluorescent cells. Then CD90.2$^{+}$ cells were sorted and within CD90.2$^{neg}$ cells, CD11c$^{+}$MHCII$^{+}$ cells were sorted. **b-h,** scRNA-seq of CD11c$^{+}$MHCII$^{+}$ cells was analysed. Clusters correspond to the UMAP in Fig. 6a. **b,** Bubble plot of selected marker genes used for cell type identification. **c,** UMAP display of AUC scores for the indicated gene signatures used for cluster identification. **d,** Distribution of cell cycle phase across clusters. **e,** Contribution of cells from each genotype to the indicated clusters as percentage. Dotted line represents fraction cells from *Clec9a$^{+/cre}$Stat1$^{fl/fl}$* mice as a percentage of total cDC2/DC3/tDC in the dataset. **f**, 3-week-old mice were injected intravenously with anti-CD45.2. Two minutes later mice were sacrificed and the indicated splenic populations analysed for CD45.2 labelling. **g,** Expression of *Stat1* in cDC1 and cDC2 clusters split by genotype. Box represents interquartile range, horizontal bar represents median, whiskers represent minimum and maximum enrichment scores. **h,** Gene set enrichment scores of the indicated gene sets. Sets that were significantly enriched in cDC1 clusters 20 or 15 are shown. Statistical analysis was performed using multiple t-tests corrected for multiple comparisons using the Holm-Šídák method, $^{*}p < 0.05$, $^{**}p < 0.01$, $^{***}p < 0.001$, $^{****}p < 0.0001$.

### Supplemental Figure 7:

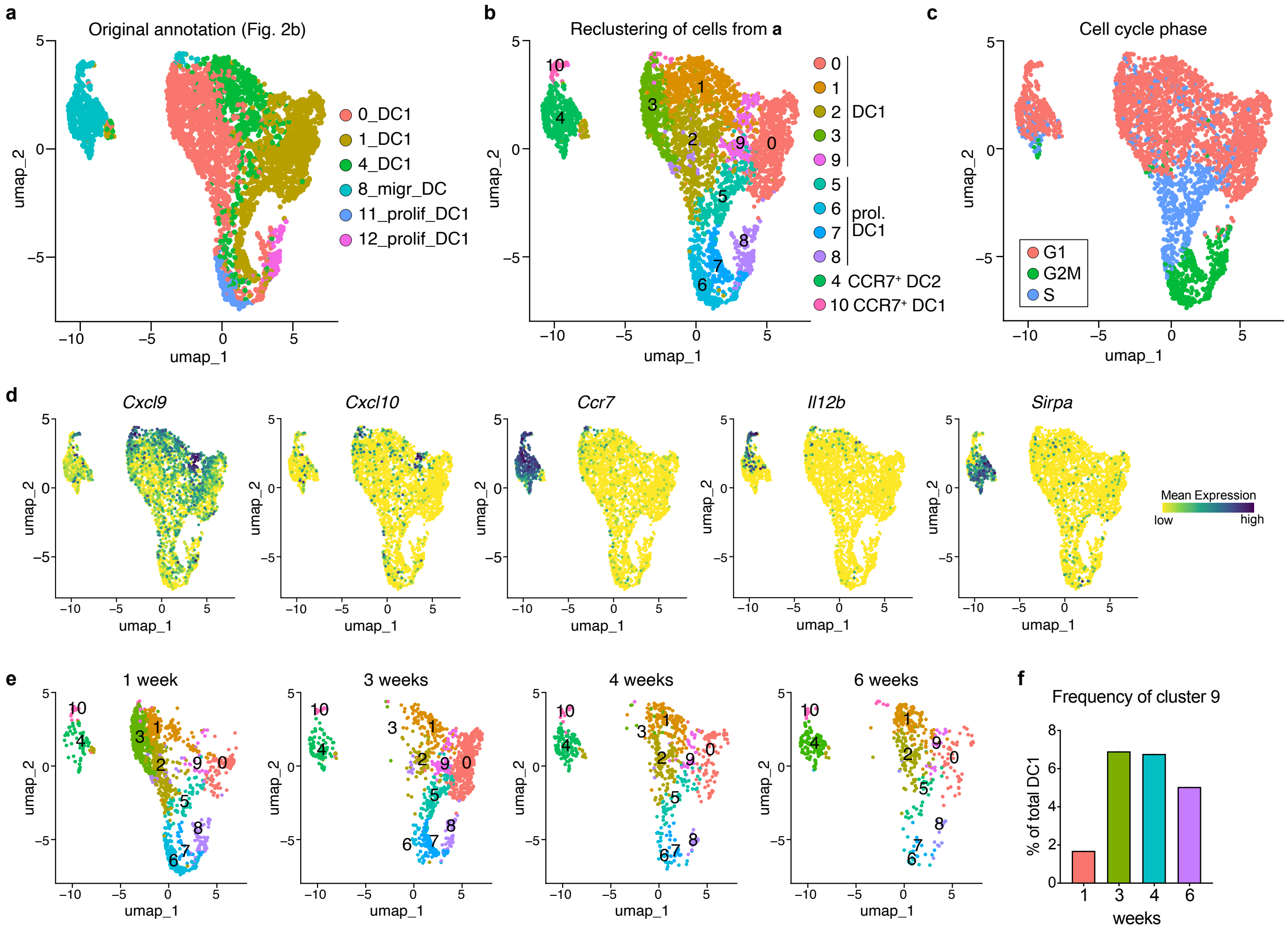

**Supplemental Figure 7: Reanalysis of isolated cDC1 and CCR7$^+$ cDC clusters from the scRNA-Seq dataset across age shown in Fig. 2**

**a-f,** cDC1 and CCR7$^+$ cDC clusters from the scRNA-seq dataset across age (Fig. 2b) were isolated and reanalysed. **a,** UMAP display of isolated cells showing their original annotation as in Fig. 2b. **b,** UMAP showing annotated clusters after renewed unsupervised clustering. **c,** UMAP displaying cell cycle phase. **d,** Expression of *Cxcl9, Cxcl10, Ccr7, Il12b,* and *Sirpa* projected on the UMAP display. **e,** UMAP split by time point. **f,** Frequency of cluster 9 as percentage of total DC1 across different time points.

### Supplemental Figure 8:

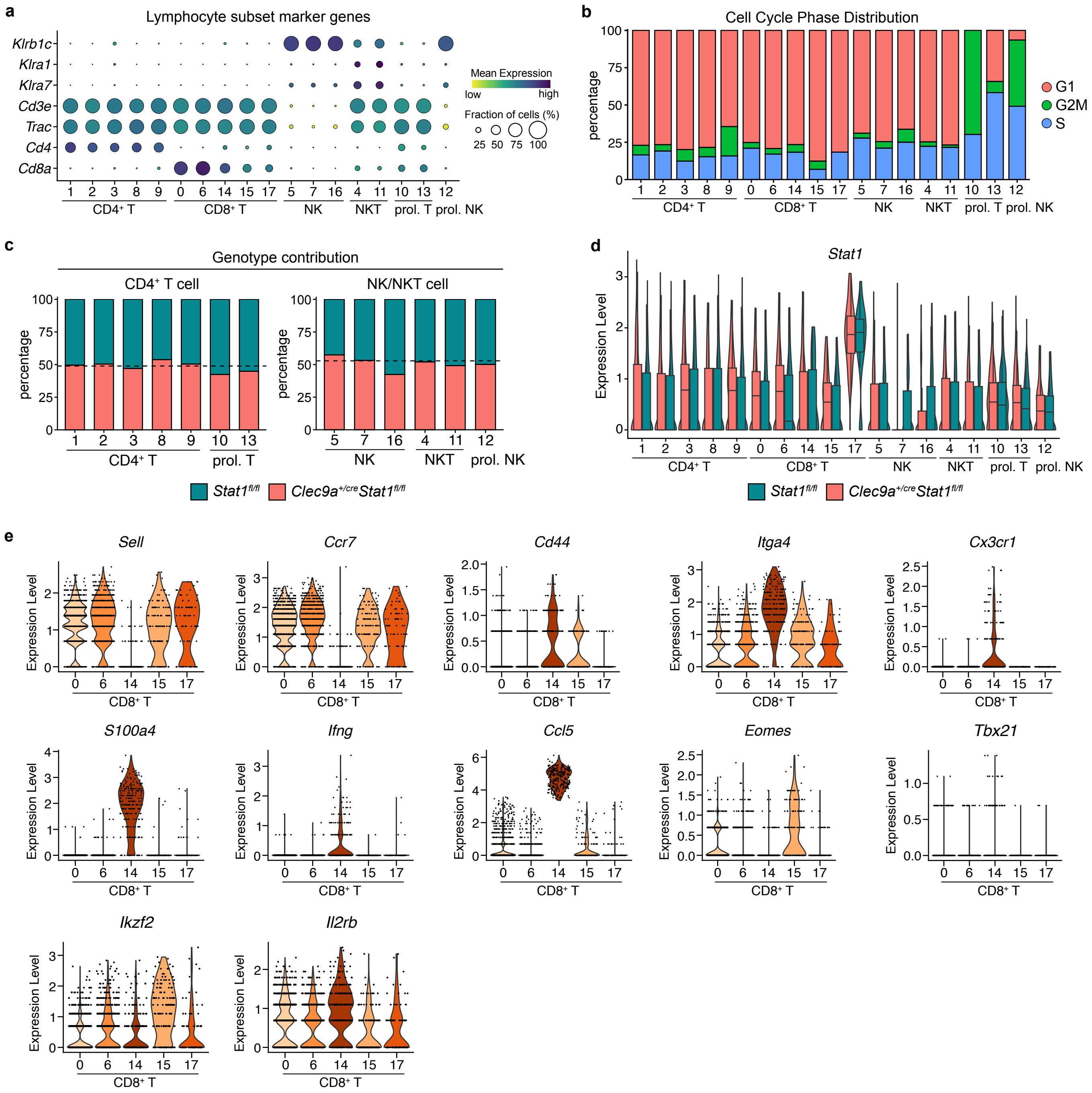

**Supplemental Figure 8: Transcriptomic analysis of spleen CD90.2$^{+}$ cells from 3-week-old *Clec9a$^{cre}$Stat1$^{fl/fl}$* mice**

**a-e,** scRNA-seq of CD90$^{+}$ cells was analysed. Clusters correspond to the UMAP in Fig. 7a. **a,** Bubble plot of selected marker genes used for cluster identification. **b,** Distribution of cell cycle phase across unsupervised clusters. **c,** Contribution of cells from each genotype to CD4$^{+}$ T cell and NK/NKT cell clusters. Dotted line represents fraction of cells from *Clec9a$^{+/cre}$Stat1$^{fl/fl}$* mice as a percentage of total T cell or NK/NKT in the dataset. **d,** Box plot showing expression of *Stat1* in the indicated clusters split by genotype. Box represents interquartile range, horizontal bar represents median, whiskers represent minimum and maximum enrichment scores. **e,** Expression of the indicated genes used for cluster identification across CD8$^{+}$ T cell clusters.

### Supplemental Figure 9:

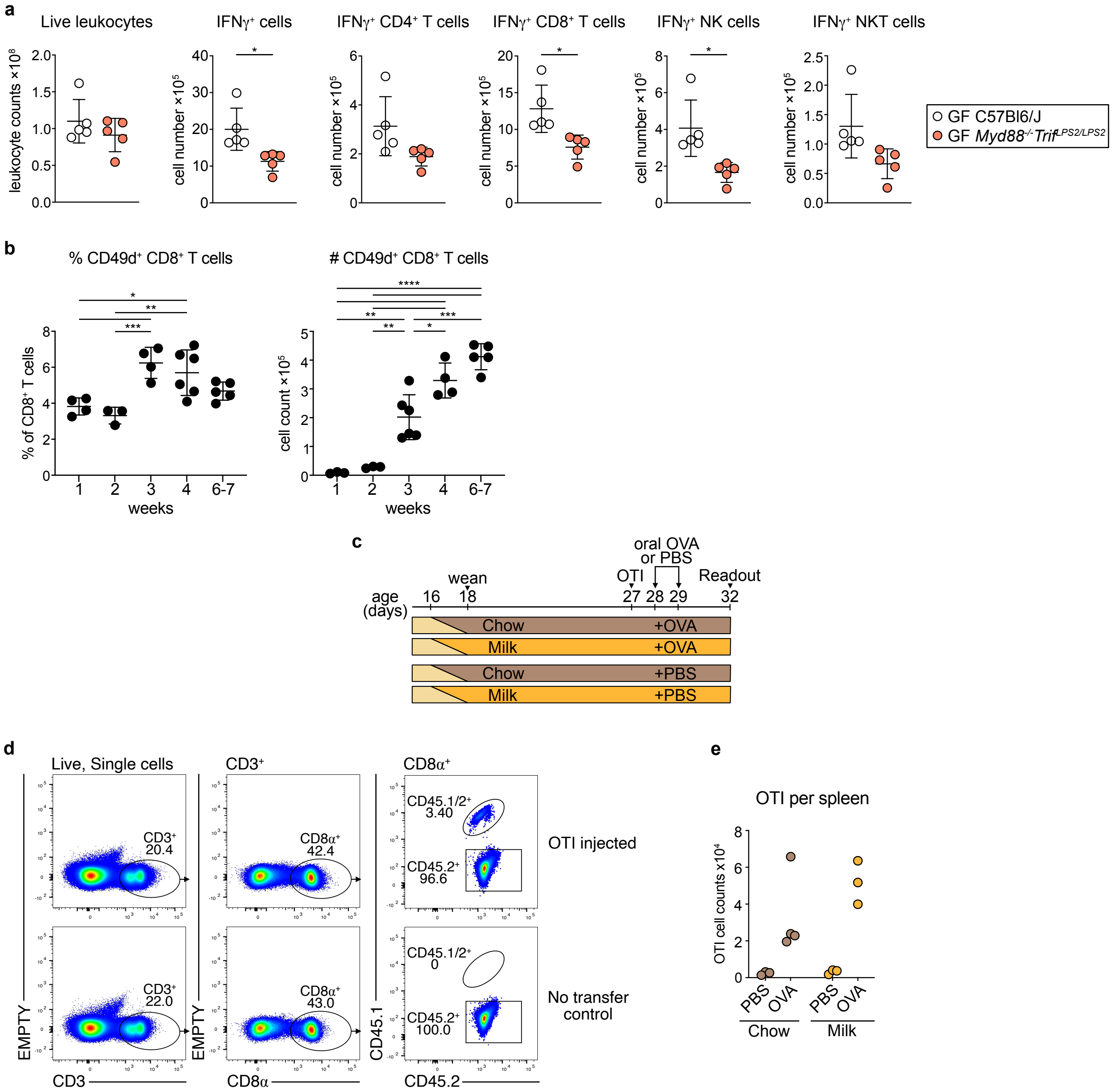

**Supplemental Figure 9:**

**a**, Total spleen cellularity, as well as the numbers of total IFNγ producing cells and the indicated of IFNγ producing lymphocyte subsets were quantified spleens from in 3-week-old GF *Myd88*$^{-/-}$*Trif*$^{LPS2/LPS2}$ or age-matched GF wild type controls. **b**, Spleens from wild type SPF mice at the indicated ages were analysed. The frequency of CD49d$^{+}$ cells as percentage of total CD8$^{+}$ T cells and numbers of splenic CD49d$^{+}$ CD8$^{+}$ T cells across age are shown. **c-e**, SPF mice were either conventionally weaned on day 18 or weaning was delayed by placing mice on formula milk from day 16 after birth while restricting access to chow. At 27 days of age, mice received $1\times10^5$ naïve OTI cell intravenously. On the next two consecutive days, mice received either oral OVA or PBS. Three days later spleens were analysed to assess OTI proliferation. **c,** Experimental scheme. **d**, Representative gating to identify OTI cells. They were identified as CD45.1/2$^{+}$ cells, distinguishing them from CD45.2$^{+}$ endogenous CD8$^{+}$ T cells. **e,** Number of recovered OTI cells in the spleen.
